# Covalent inhibitors of the RAS binding domain of PI3Kα impair tumor growth driven by RAS and HER2

**DOI:** 10.1101/2024.12.17.629001

**Authors:** Joseph E. Klebba, Nilotpal Roy, Steffen M. Bernard, Stephanie Grabow, Melissa A. Hoffman, Hui Miao, Junko Tamiya, Jinwei Wang, Cynthia Berry, Antonio Esparza-Oros, Richard Lin, Yongsheng Liu, Marie Pariollaud, Holly Parker, Igor Mochalkin, Sareena Rana, Aaron N. Snead, Eric J. Walton, Taylor E. Wyrick, Erick Aitichson, Karl Bedke, Jacyln C. Brannon, Joel M. Chick, Kenneth Hee, Benjamin D. Horning, Mohamed Ismail, Kelsey N. Lamb, Wei Lin, Justine Metzger, Martha K. Pastuszka, Jonathan Pollock, John J. Sigler, Mona Tomaschko, Eileen Tran, Todd M. Kinsella, Miriam Molina-Arcas, Gabriel M. Simon, David S. Weinstein, Julian Downward, Matthew P. Patricelli

## Abstract

Genetic disruption of the RAS binding domain (RBD) of PI 3-kinase (PI3K) prevents the growth of mutant RAS driven tumors in mice and does not impact PI3K’s role in insulin mediated control of glucose homeostasis. Selectively blocking the RAS-PI3K interaction may represent an attractive strategy for treating RAS-dependent cancers as it would avoid the toxicity associated with inhibitors of PI3K lipid kinase activity such as alpelisib. Here we report compounds that bind covalently to cysteine 242 in the RBD of PI3K p110α and block the ability of RAS to activate PI3K activity. These inhibitors have a profound impact on the growth of RAS mutant and also HER2 over-expressing tumors, particularly when combined with other inhibitors of the RAS/MAPK pathway, without causing hyperglycemia.

## Introduction

*RAS* oncogenes are mutationally activated in 20% of human cancers, with RAS proteins activating both the MAPK and PI3K pathways (*1-3*). As each of these pathways has oncogenic potential, simultaneous activation, as occurs in RAS driven cancers, generates aggressive disease. In RAS driven cell and animal models, dual inhibition of the MAPK and PI3K pathways has shown superior efficacy relative to targeting the individual pathways (*4*), however, dose limiting toxicities in humans have prevented this combination strategy from finding clinical success. While physiological activation of the MAPK pathway is RAS dependent, the interaction between RAS and the catalytic subunit of PI3Kα, p110α, serves as an amplifier but not a primary activator of this pathway, and is less important in normal cellular regulation compared to in cancer (*5, 6*).

Genetic disruption of the RAS-p110α interaction blocks progression of mutant RAS induced lung and skin tumors in mice, highlighting its importance in RAS-driven cancer cells (*6, 7*). Although the interaction of RAS with p110α was disrupted in all tissues of these mice, there was no identifiable impact on fitness of adult animals, indicating that in healthy cells, RAS dependent amplification of PI3K signaling is expendable and RAS independent activation of PI3K by upstream signaling factors is sufficient for maintaining physiological homeostasis. Unfortunately, targeting the PI3K pathway with inhibitors of the catalytic activity of p110α have had limited clinical success due to on-target dose-limiting toxicities, most commonly hyperglycemia and rash, caused by inhibition of both RAS-dependent and -independent PI3K signaling (*8, 9*). Insulin regulation of glucose homeostasis is severely disrupted by p110α lipid kinase inhibitors, including alpelisib which is approved for the treatment of breast cancer (*10*).

Here we describe small molecules that covalently ligate cysteine 242, adjacent to the RAS binding interface of p110α, blocking its ability to interact with RAS proteins. These compounds inhibit the growth of RAS mutant tumors, particularly when combined with MEK or KRAS inhibitors. Surprisingly, these compounds also inhibited apparently RAS independent activation of PI3K signalling in HER2 over-expressing cells, suggesting that the RBD may play a non-canonical role in this setting. The molecules were well tolerated, did not impact normal blood glucose regulation and are promising candidates for cancer therapy. A related molecule is entering Phase I clinical trials in humans.

## Results

### Identification of compounds that bind covalently to the RAS binding domain of p110α

RAS proteins associate with their effector proteins through electrostatic as well as hydrophobic interactions. Mutational studies have identified key residues, T208 and K227, within the RAS binding domain (RBD) region of p110α, which facilitate the interaction with RAS upon receptor tyrosine kinase driven recruitment of PI3Kα (**Fig. 1A**). Using mass-spectrometry-based chemoproteomics (*11*) we were able to detect 21 cysteines on p110α, including cysteine 242, which is exclusive to p110α, adjacent to the RBD and proximal to T208 and K227. Additionally, comparison of unbound p110α to p110γ bound to HRAS (*12*) indicates that C242, analogous to P262 of p110γ, sits on an α-helix that undergoes a conformational rearrangement upon RAS binding (**Fig. 1A**). This led us to search for covalent ligands of C242 of p110α as potential blockers of the interaction between these two proteins.

**Figure 1:**
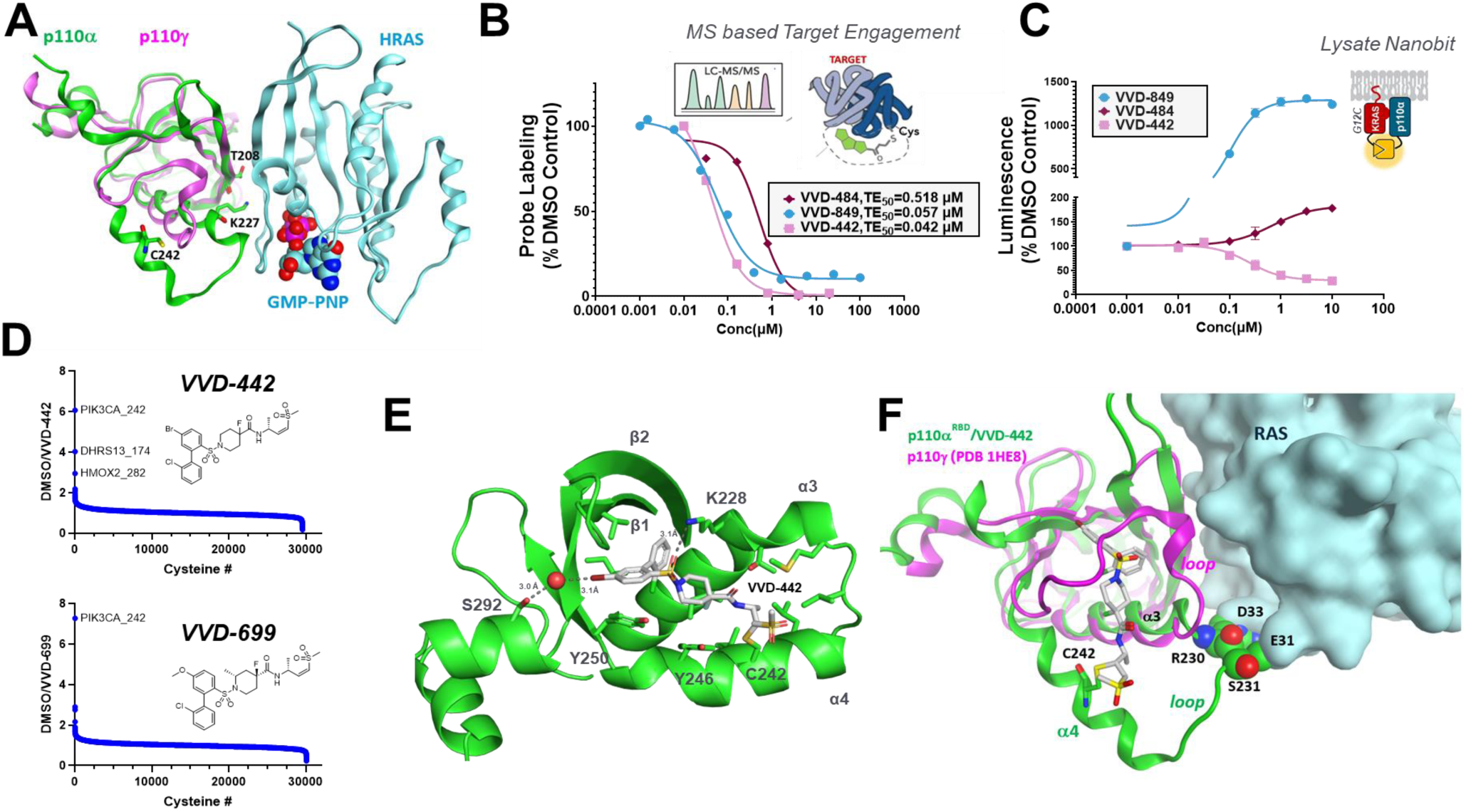
Identification of compounds that bind covalently to the RAS binding domain of p110α. (A) Ribbon representation of the p110γ/RAS/GMPPNP crystal structure (PDB code 1HE8) with the superimposed coordinates of p110α(6V07). Residues T208 and K227 of p110α are located at the interface of the RBD region and RAS and facilitate interactions between the proteins. Residue C242 is located in the α4-helix. In the p110γ/RAS crystal structure, the α4-helix undergoes conformational changes upon RAS binding highlighting high flexibility of the region. RAS, p110γ, and p110α are shown in light blue, magenta and green colors, respectively. GMPPNP is represented as a space-filling model. (B) LC-MS/MS based target engagement (TE) was calculated by comparing the probe-labeled PIK3CA_C242 areas under the curve (AUCs) of compound treated to control jurkat cell lysates. (C) VVD-442, VVD-849, VVD-484 were evaluated for their ability to impact the interaction between p110α and KRAS^G12C^ using the Nanobit protein-protein interaction assay in HEK293T cell lysates treated with compound for 1 hour. (D) Global proteomics selectivity of VVD-442 and VVD-699. Compound selectivity relative to ∼30,000 cysteine sites was measured at ∼25-fold over IC_50_s (2 µM, 2 h live cell treatment) using TMT quantification. (E) Crystal structure of the VVD-442-RBD complex. The compound rests in a hydrophobic cleft formed by α3 and α4 and β-strands 1 and 2. (F) Model of the RAS/p110α/VVD-442 interaction showing steric clashes between residues R230 and S231 of p110α and residues E31 and D33 of RAS. Additionally, in the p110γ/RAS X-ray structure, a flexible loop (A253-Q268) undergoes notable conformational changes upon RAS binding. The p110γ and p110α RBD subunits are shown in magenta and green colors, respectively, while RAS is depicted in surface representation. VVD-442 is shown as a stick model.

Towards this end, we used our targeted chemoproteomics screening platform to identify compounds representing multiple chemotypes that selectively formed a covalent bond with C242 (**Fig 1B**). Medicinal chemistry optimization led to the generation of potent and selective C242 engagers with diverse structural features (**Fig. S1A**). To understand the consequence of C242 ligation, we employed a NanoBiT assay in HEK293T cell lysates to measure the protein-protein interaction between KRAS^G12C^ and p110α (*13*). This assay revealed that compounds binding to p110α C242 fall into three different classes: blockers of the interaction, promoters of the interaction (glues) or compounds lacking effect on the interaction (silent ligands) (**Fig 1C, Fig. S1A, Table S1**). As our goal was to develop a small molecule that can disrupt the RAS-p110α interaction, we focus in this study on a number of blockers: early tool compounds VVD-442(**Fig. 1C**) and VVD-699, and the further optimized molecules VVD-844 and VVD-579 (**Fig. S1A-B, Fig. S2A, Table S1**). Global proteomic profiling of ∼30,000 cysteine residues at ∼25-fold over compound IC_50_s revealed a favorable selectivity profile for VVD-442, which improved with VVD-699 (**Fig. 1D**).

To understand how our covalent ligands for Cys242 block the interaction between RAS and p110α, we determined the X-ray co-crystal structure of VVD-442 covalently bound to the RBD of p110α at 2.83 Å resolution (**Fig 1E, Fig. S1C** and **Table S2**). The compound, covalently bound to Cys242 lies in a hydrophobic groove between α-helices 3 and 4. The central sulfonamide forms a hydrogen bond with the terminal amide of Lys228 and the bromine forms a halogen bond (or σ-hole interaction) with a water positioned by Ser292. Comparison of the apo (PDB ID 6VO7) and liganded structures shows a 2.8 Å shift in the N-terminus of helix 4 and a 2.4 Å shift in the position of the Tyr246 side chain (**Fig. S1D**) (*14*). Each of these movements is required for the molecule to form the pocket defined by helices 3 and 4. VVD-442 binds on the opposite side of β-strands 1 and 2 of the RBD, the site of the primary interaction with RAS, suggesting that the molecule does not block RAS-p110α binding through direct steric occlusion. Modeling the p110α/VVD-442 complex with RAS, using the crystal structure of p110γ bound to RAS as a template (PDB ID 1HE8), suggests that the conformation of the (α3-α4) loop, observed in the inhibitor-bound state, is incompatible with the conformation of the switch-I region of RAS (**Fig. 1F**). Specifically, residues Arg230 and Ser231 of p110α sterically clash with residues Glu31 and Asp33 of RAS, disrupting both the electrostatic and structural complementarity required for complex formation. This model is consistent with experimental data showing that binding of VVD-442 to p110α disrupts the RAS-p110α interaction.

### Cellular characterization of the RAS-p110α blocker VVD-699

To confirm that VVD-699 disrupts the RAS-p110α interaction in cells, we initially focused on KRAS mutant H358 lung cancer cells, which have been shown to rely on KRAS^G12C^ for maximal PI3K activation (*15*). VVD-699 engaged p110α in H358 cell lysates based on competition with a biotinylated probe **(Fig. S2B** for assay design and structure of probe) and, with comparable potency, inhibited downstream AKT phosphorylation in cells (**Fig. 2A**). Importantly, VVD-699 had no impact on p110α lipid kinase activity in a biochemical assay, supporting that cellular pAKT inhibition is not due to direct inhibition of kinase activity (**Fig. 2B**). VVD-699 also inhibited pAKT in KRAS^G12S^ A549 cells, and this inhibition was lost in isogenic p110α C242S knock-in A549 cells (**Fig. 2C**). To further confirm VVD-699 is blocking RAS promoted p110α activity, we utilized mouse embryonic fibroblasts (MEFs) which have been engineered to express an HRAS-G12V-ER fusion protein, allowing for rapid inducible activation of RAS upon addition of 4-hydroxytamoxifen (4HT) (*6*). p110α-WT MEFs pre-treated with VVD-699 prior to 4HT addition also showed near complete blockage of HRAS-G12V driven activation of PI3K (**Fig. 2D**). In MEFs expressing p110α with blocking mutations in the RBD (T208D and K227A), the ability of inducible RAS to activate PI3K activity was greatly impaired, as previously shown (*6*).

**Figure 2:**
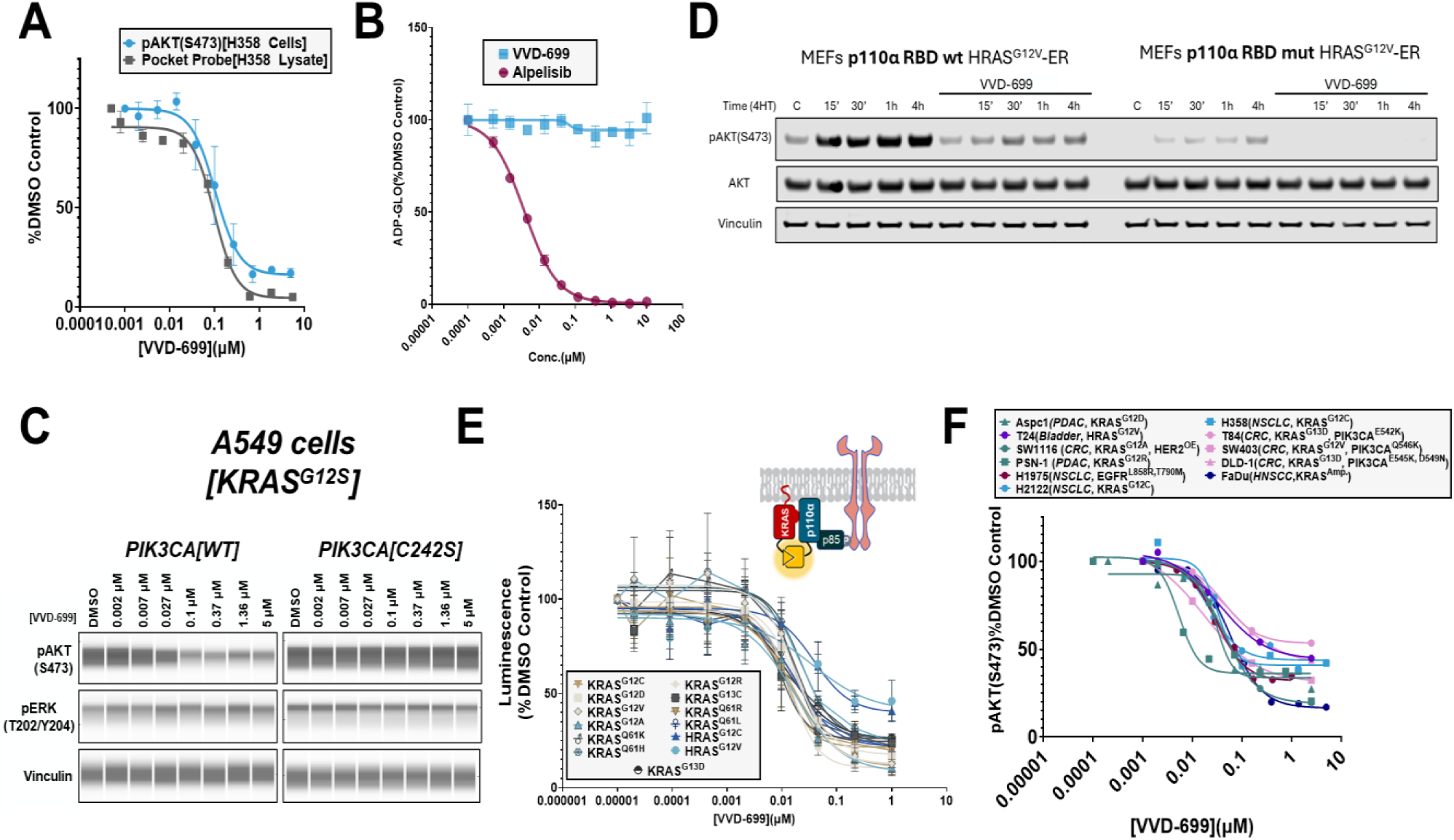
VVD-699 blocks the RAS-p110α interaction in cells and inhibits PI3Kα activation. (A) H358 cell lysates treated with VVD-699 for 2 hours to measure p110α-C242 target engagement through pocket probe analysis and H358 cells treated with VVD-699 for 30 minutes to measure pAKT(S473) through HTRF. (B) p110α kinase activity assay performed in the presence of VVD-699 or alpelisib. (C) A549 cells, PIK3CA WT or C242S, treated with VVD-699 for 2 hours to measure activation of the PI3K/AKT and MAPK pathways through western blot analysis. (D) p110α^WT^ or p110α^mut^ ^(T208D/K227A)^ MEFs were pre-treated with DMSO or VVD-699 followed by addition of tamoxifen to induce expression of HRAS-G12V. Samples were collected at the indicated timepoints and subjected to western blot analysis. (E) VVD-699 was screened for its ability to disrupt the interaction between p110α and various RAS mutants using the Nanobit protein-protein interaction assay in H358 cells. (F) VVD-699 was screened across numerous cell lines and activation of PI3K was measured through assessment of pAKT(S473) through HTRF or western blot analysis.

We next evaluated the ability of VVD-699 to impact p110α interactions with RAS and the impact on downstream signaling across a variety of mutations that lead to PI3K-AKT pathway activation. In a live cell NanoBiT assay, VVD-699 blocked the interaction of all tested KRAS and HRAS mutants with p110α with broadly similar potency and magnitude of disruption (**Fig. 2E**). In addition, a p110α-helical domain mutant showed similar behavior (**Fig. S2C**). Consistent with the NanoBiT findings, VVD-699 inhibited pAKT in numerous cancer cell lines harboring various RAS mutations, as well as cell lines with co-mutations to KRAS and p110α, with varying levels of maximal inhibition, but clear activity in all cases (**Fig. 2F)**. Collectively these findings show that VVD-699 is able to disrupt the interaction of p110α and RAS through covalent liganding of C242 and block downstream pathway activation across a wide range of cancer relevant RAS and p110α mutations.

### VVD-699 inhibits HER2 activity through an H/K/N-RAS independent mechanism

Despite the clear impact of VVD-699 on p110α-RAS interaction and inhibition of downstream signaling, we only observed a modest impact on cell growth of KRAS mutant cell lines, including those with PIK3CA co-mutation, in proliferation assays carried out in anchorage independent settings (**Fig. S2D**). Surprisingly, however, we found that cell lines harboring amplification of HER2 were particularly sensitive to VVD-699, exhibiting deeper inhibition of pAKT than in RAS mutant settings, and clear inhibition of cell growth in culture (**Fig. 3A-B**). To explore the mechanism of this robust HER2 signaling inhibition, we evaluated the glue and silent ligands described above that potently engage p110α C242 but differentially impact the p110α-RAS interaction (**Fig. 1C**). In the EGFR mutant cell line H1975, which we have previously shown to be partly dependent on the interaction of RAS with p110α (*16*), as well as the KRAS amplified (KRAS^amp^) cell line FaDu (*17*), the compounds had impacts on pAKT signaling (**Fig. 3C**) consistent with their behavior in the RAS-p110α NanoBiT assay (**Fig. 1C**). However, in the HER2 overexpressing cell line N87 (*18*), compound performance was disconnected from the ability to disrupt the RAS-p110α interaction. Specifically, both VVD-699, a blocker, and VVD-484, a silent ligand, achieve near full pAKT(S473) inhibition, while VVD-849, a glue, did not induce pAKT(S473) signaling in N87 cells, but exhibited partial pAKT(S473) inhibition, suggesting a mixed impact of this ligand. These results support that the activity of p110α C242 ligands in HER2 signaling is occurring at least predominantly through a RAS-independent mechanism. To investigate this further, we treated KRAS mutant and HER2 overexpressing cells with the multi-RAS (ON) inhibitor RMC-6236 (*19*). In both settings, RMC-6236 was highly effective at inhibiting the MAPK pathway (**Fig. S3A-B**). In KRAS mutant cells, we find that RMC-6236 and VVD-699 both achieve partial inhibition of pAKT(S473) with comparable activity in most cell lines(**Fig. 3D**). However, in HER2 overexpressing cells, RMC-6236’s ability to inhibit PI3K activation is highly variable, while VVD-699 provided near full PI3K inhibition (**Fig. 3E**). Comparison of pAKT(S473) inhibition by RMC-6236 and VVD-699 in KRAS^mut^ cells resulted in good agreement through linear regression analysis (R^2^=0.64) while an analogous analysis in HER2 over-expressing (HER2^OE^) cells displayed minimal agreement (R^2^=0.12) (**Fig. S3C**). To further explore the possibility that VVD-699 is functioning in a RAS-independent manner in cells with high levels of HER2, we overexpressed p110α-RBD, which functions as a dominant negative inhibitor of the RAS-p110α interaction (*16*), in A431 (EGFR^OE^) and N87 (HER2^OE^) cells. This resulted in dramatic inhibition of pAKT(S473) in A431 cells but only a minor impact in N87 cells. Importantly, the addition of VVD-699 to N87 cells overexpressing p110α-RBD led to near complete inhibition of pAKT(S473) (**Fig. S3D**). Finally, we explored the possibility that p110α-C242 ligands might prevent HER2 signaling through impacts on non-canonical RAS isoforms that are not inhibited by RMC-6236 such as MRAS and RRAS. We utilized our live cell Nanobit system and found that, although the luminescence signal varied between the isoforms, introduction of the RBD mutations readily disrupts the interaction between multiple RAS related proteins and p110α (**Fig. 3F**). Interestingly, we found that none of the VVD ligands, including VVD-699, were strong blockers of the interaction between p110α and these alternate RAS isoforms. In fact, VVD-484, which shows strong pAKT(S473) inhibition in N87 cells, functions either as a silent ligand or an activator when p110α is paired with the alternate RAS isoforms (**Fig. 3G**). In sum, these data indicate that the activity of VVD-699 in the HER2 OE setting is driven by a distinct and receptor-specific mechanism that is orthogonal to the molecule’s ability to disrupt RAS-p110α interactions.

**Figure 3:**
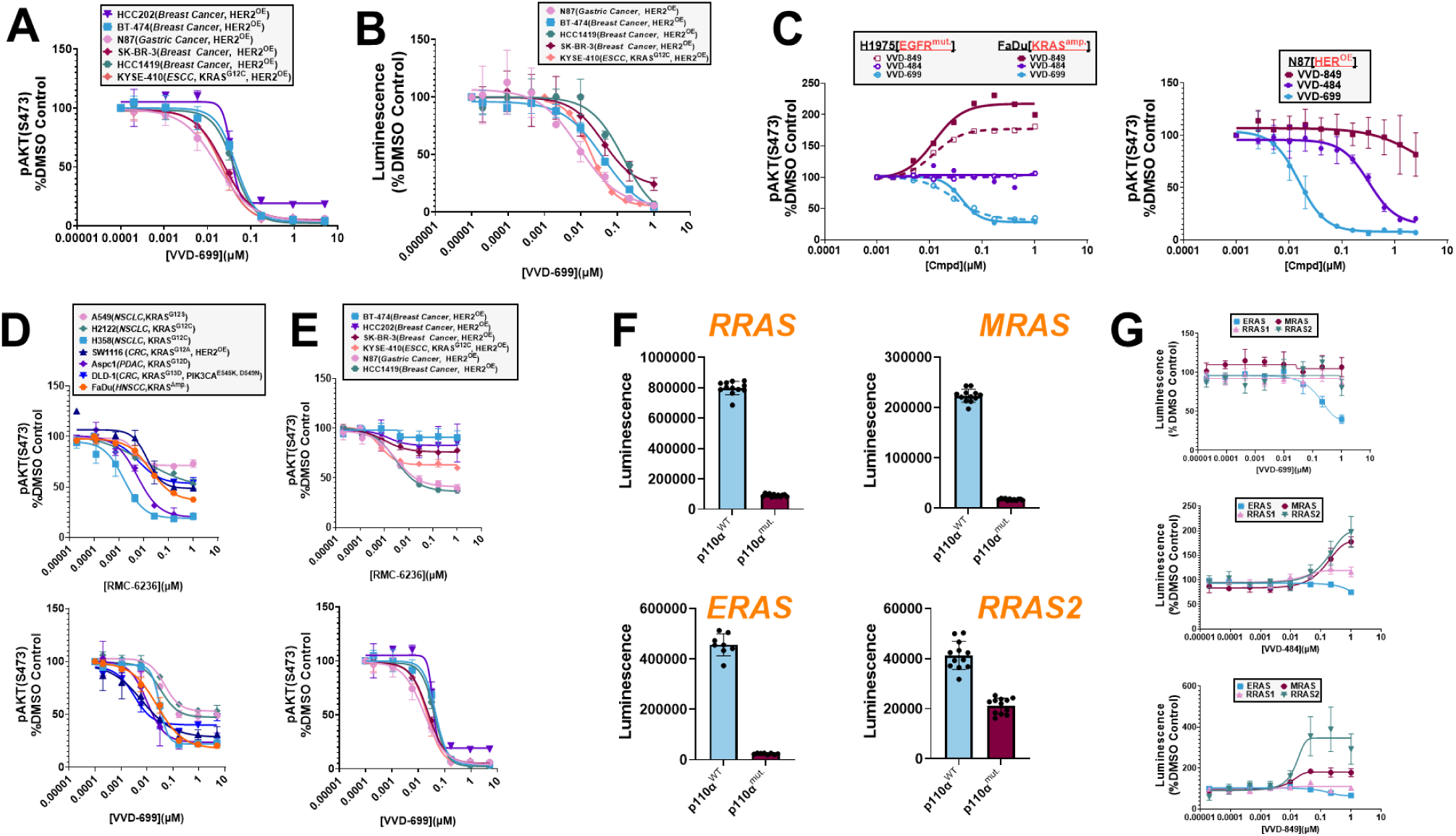
HER2 overexpressing cells show unique sensitivity to VVD-699. (A) HER2 overexpressing cells treated with VVD-699 for 2 hours followed by analysis of pAKT(S473) through HTRF. (B) HER2 overexpressing cells treated with VVD-699 every 3 days for either 6 or 9 days, followed by assessment for cell abundance using Cell Titer Glo. (C) H1975(EGFR_mut_) or FaDu(KRAS^amp^) cells, driven RAS activity, or N87 cells, driven by HER2 overexpression, were treated with either VVD-849, VVD-484 or VVD-699 for 2 hours, followed by analysis of pAKT(S473) through HTRF. (D/E) Cells driven by RAS activity (D) or HER2 overexpression (E) were treated with RMC-6236 or VVD-699, followed by analysis of pAKT(S473) through HTRF. (F/G) The interaction between the identified RAS isoforms and p110α^WT^ or p110α^mut.^ using the Nanobit protein-protein interaction assay in H358 cells (F). Using this same assay format, the ability of VVD-849, VVD-484 or VVD-699 to impact the interaction between the identified RAS related isoforms and p110α^WT^ was determined (G).

### VVD-699 Inhibits Tumor Growth Without Impacting Glucose Homeostasis

While we did not observe broad impact on cell growth with our p110α-RAS blockers *in vitro*, we speculated that the biological importance of the p110α-RAS interaction may be more pronounced *in vivo* based on prior publications using numerous mouse models (*6, 7, 16*). To confirm that VVD-699 is active *in vivo*, mice bearing FaDu xenografts (KRAS^amp^), were given a single 100 mg/kg oral dose of VVD-699. 6 hours post-dose, near maximal target engagement as well as partial inhibition of pAKT(S473) were observed **(Fig. 4A).** Mice carrying FaDu xenografts were dosed orally for 3 days with a range of VVD-699 dose levels using a twice daily (BID) schedule, followed by assessment of tumor target engagement 6 hours post final dose. Near maximal target engagement was seen with a 30 mg/kg BID dosing schedule, which was then used for subsequent efficacy studies (**Fig. 4B**). VVD-699 induced modest tumor growth inhibition comparable to a 19 mg/kg oral dosing of alpelisib, a dose that mimics the clinical exposure of alpelisib (**Fig. 4C**). Importantly, the clinical exposure of alpelisib has been optimized to minimize the risk of inducing hyperglycemia, a known on-target dose limiting toxicity, in patients (*8, 10*). All treatment regimens were well-tolerated (**Fig. S4A**). To evaluate the impact of VVD-699 on glucose handling and in turn insulin production, we performed a time-course measuring blood glucose as well as terminal plasma insulin in mice following 3 days of dosing with either alpelisib or VVD-699. Mice dosed orally with alpelisib at 19 mg/kg QD showed a modest impact on glucose handling, while the impact on insulin production was more pronounced. However, the impact on both blood glucose levels and insulin production was dramatically increased by increasing the dose to 50 mg/kg (**Fig. 4D, Fig. S4B**). Thus, an oral dose of alpelisib in mice that is similar to the clinically approved dose in humans is only slightly below the level that causes major disruption of glucose homeostasis. Conversely, neither 30 nor 100 mg/kg of VVD-699, both of which achieved near maximal target engagement, induced a significant impact on either blood glucose or plasma insulin levels (**Fig. 4D, Fig. S4B**). These results highlight that blocking the RAS-p110α interaction without impacting intrinsic kinase activity (**Fig. 2B**) avoids disruption of glucose control.

**Figure 4:**
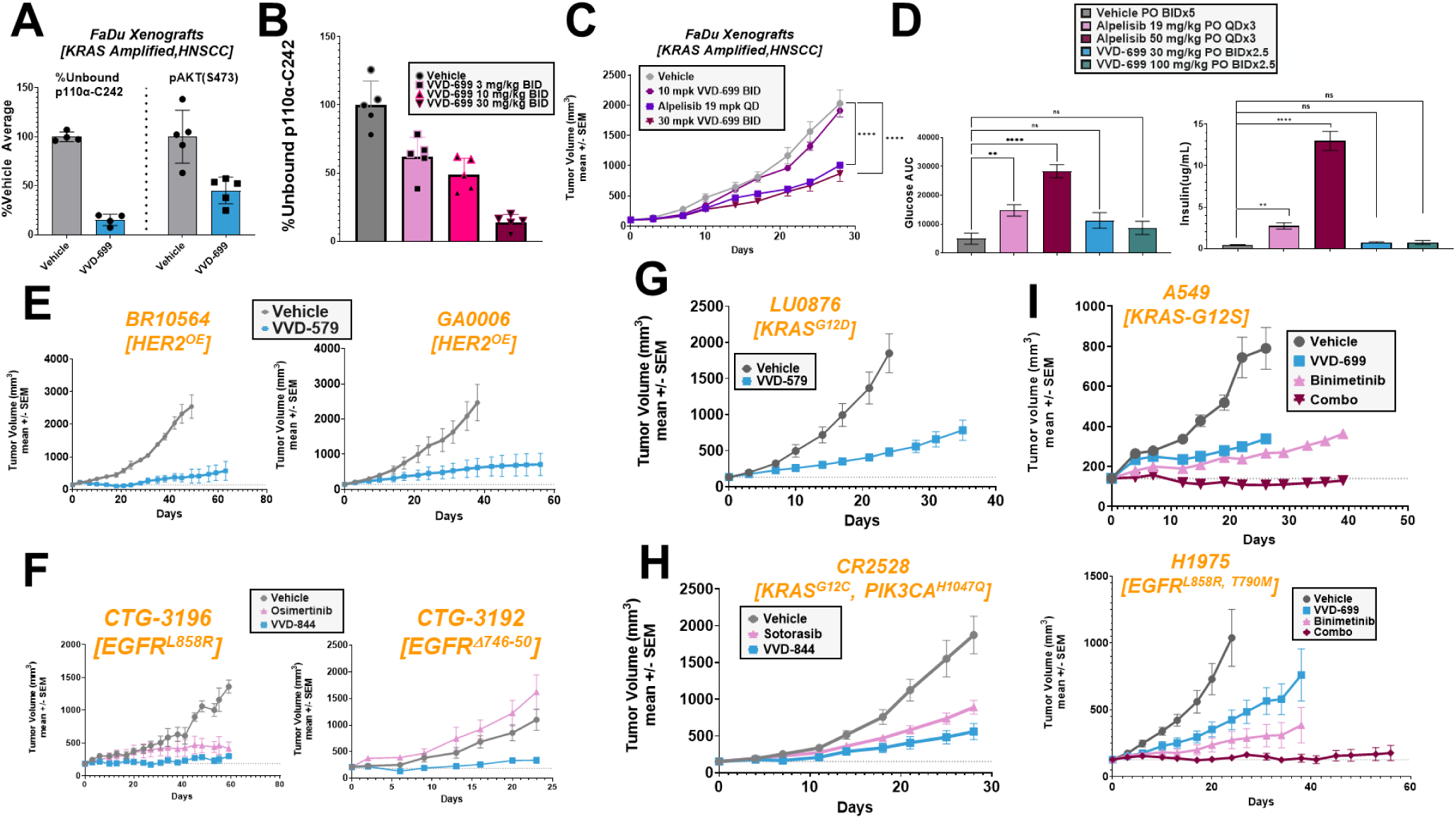
Disruption of the RAS-PI3K interaction slows tumor growth. (A) Mice bearing FaDu(KRAS^Amp^) xenografts were given an oral 100 mg/kg dose of VVD-699. 6 hours post-dose, tumor samples were collected and p110α-C242 target engagement was measured via mass spectrometry and pAKT(S473) levels were measured via western blotting. (B) Mice bearing FaDu(KRAS^Amp^) xenografts were dosed orally with 3, 30 or 30 mg/kg dose of VVD-699 for 3 days. 6 hours post-dose, tumor samples were collected and p110α-C242 target engagement was measured through mass spectrometry. (C) Anti-tumor efficacy of VVD-699 in FaDu(KRAS^amp^) xenografts. Data are shown as mean ± SEM; n=10 animals/group. Mice were dosed orally with vehicle, 10 mg/kg VVD-699 BID, 30 mg/kg VVD-699 BID or 19 mpk alpelisib QD. (D) Mice were administered the identified doses of alpelisib or VVD-699 for 3 days, followed by serial measurement of blood glucose levels for 4 hours post-last dose. Blood glucose AUC was then calculated for each treatment group. Additionally, at the conclusion of the time course, plasma was collected and insulin level were measured. (E) Anti-tumor efficacy of VVD-579 in HER2^OE^ PDX models BR10564 and GA0006. Data are shown as mean ± SEM; n=3 animals/group. Mice were dosed orally BID with 30 mg/kg VVD-579. (F) Anti-tumor efficacy of VVD-844 in EGFR^mut^ PDX models CTG-3196 and CTG-3192. Data are shown as mean ± SEM; n=3 animals/group. Mice were dosed orally BID with 10 mg/kg VVD-172844 or orally QD with 25 mg/kg osimertinib. (G) Anti-tumor efficacy of VVD-579 in KRAS^G12D^ PDX model LU0876. Data are shown as mean ± SEM; n=3 animals/group. Mice were dosed orally BID with 30 mg/kg VVD-579. (H) Anti-tumor efficacy of VVD-844 in KRAS^G12^, PIK3CA^H1047Q^ PDX model CR2528. Data are shown as mean ± SEM; n=5 animals/group. Mice were dosed orally BID with 10 mg/kg VVD-172844 or orally QD with 100 mg/kg sotorasib. (I) Anti-tumor efficacy of VVD-699, Binimetinib, or a combination of both in A549(KRAS^G12S^)(Top) or H1975(EGFR^L858R, T790M^)(Bottom) xenografts. Data are shown as mean ± SEM; n=10 animals/group. Mice were dosed orally BID with 30 mg/kg VVD-699, QD with 30 mg/kg binimetinib or a combination of both

To further characterize the breadth of efficacy of p110α C242 ligands, we evaluated VVD-699, and tool compounds representing alternative chemotypes with similar activity(**Fig. S2A, Table S1**), across a variety of tumor models. Strong growth inhibition was seen in patient derived xenograft (PDX) models that overexpress HER2 (**Fig. 4E**), consistent with our *in vitro* signaling results. Additionally, single agent activity was found in EGFR^mut^ PDX models, including models derived from patients that developed resistance to Osimertinib (**Fig 4F**). We also identified sensitivity to these compounds in numerous KRAS^mut^ PDX models, including models carrying the two most common KRAS mutations, G12D (**Fig. 4G**) and G12V (**Fig. S4C**). Disruption of the RAS-p110α interaction also suppressed growth of a PDX model carrying co-mutation to both KRAS and p110α, in this case providing comparable activity to direct targeting of KRAS^G12C^ (**Fig. 4H**). These data highlight the importance of the RAS-p110α interaction across a variety of backgrounds; however, the lack of complete tumor growth inhibition supports prior observations that targeting individual RAS effector pathways results in limited efficacy (*20*). With this in mind, we explored disruption of the RAS-p110α interaction in combination with a MEK inhibitor, to simultaneously inhibit the PI3K and MAPK pathways. A549 (KRAS^G12S^) cells treated *in vitro* with VVD-699 and binimetinib showed inhibition of both the PI3K/AKT and MAPK pathways, supporting evaluation of this strategy (**Fig. S4D**). We next tested this combination *in vivo* using A549 and H1975 xenografts (**Fig. 4I**). In both models, the combination treatment generated a significantly improved response relative to either single agent, with durable tumor stasis observed. Interestingly, although H1975 was far more responsive than A549 to VVD-699 in 3D proliferation assays (**Fig. S2D**), similar efficacy is seen *in vivo*, highlighting the challenge of assessing sensitivity to this mechanism of action *in vitro*. Lastly, we evaluated this combination in KRAS^mut^ PDX models, finding clear combination benefit relative to the individual therapies across models carrying varying mutations to KRAS (**Supp. Fig. 4E**).

### VVD-699 sensitizes tumors to KRAS^G12C^ Inhibition

Although patients initially respond to KRAS^G12C^ inhibitors, tumors quickly develop resistance to the treatment (*21*). Post-treatment analysis indicates that this occurs through both genetic and non-genetic mechanisms of resistance, nearly all of which lead to reactivation of the RAS signaling pathways (*22*). We initially focused on H2122 cells, previously characterized as intrinsically partially resistant to KRAS^G12C^ inhibition (*23*). When these cells are treated with sotorasib for 2 hours, both the MAPK and PI3K/AKT pathways are effectively suppressed, highlighting the role of KRAS^G12C^ in activation of these pathways (***Fig. 5A***). However, after 24 hours of sotorasib treatment, while MAPK pathway inhibition is maintained, signaling through the PI3K/AKT pathway is restored. In contrast, VVD-699 maintains pAKT inhibition in H2122 cells at both 2 and 24 hours (**Fig. S5A**). Combination treatment with VVD-699 and sotorasib suppressed both PI3K/AKT and ERK signaling at 2 and 24 hours, indicating that this combination can be used to more durably suppress the PI3K/AKT and MAPK pathways in KRAS^G12C^ driven cancers (**Fig. 5B, Fig. S5B**). As wild-type RAS isoforms are known to support signaling in KRAS^mut^ cancers (*24*), we evaluated the role of wild-type H/NRAS in H2122 cells, which are homozygous for KRAS^G12C^. CRISPR knockdown of H/NRAS in H2122 cells led to a decrease in pAKT(S473), but not pERK(T202/Y204) (**Fig. S5C**). These results indicate that while activation of the MAPK pathway is dominantly controlled by KRAS-G12C in H2122 cells, activation of the PI3K/AKT pathway is more complex and has contributions from WT-RAS isoforms. Additionally, as wild type H/NRAS play a role in the basal signaling network of H2122 cells, we speculated that they may also play a role in driving the rebound in PI3K/AKT signaling seen with prolonged inactivation of KRAS^G12C^. In support of this hypothesis, the multi-RAS (ON) inhibitor RMC-6236 maintained pAKT inhibition at 24 hours treatment, albeit it with decreased potency (**Fig. 5C**).

**Figure 5:**
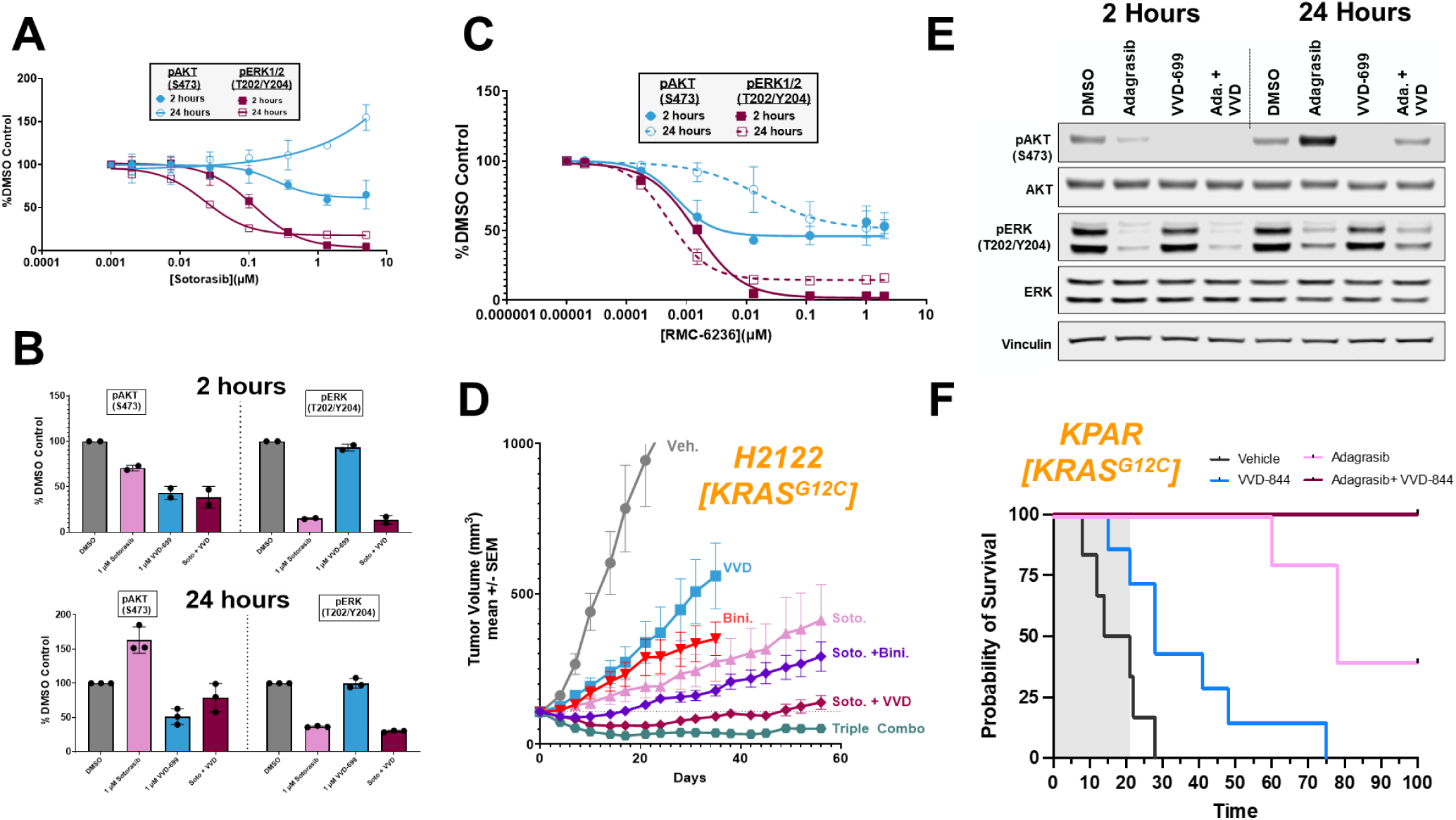
VVD-699 provides benefit with used in combination with KRAS-G12Ci. (A) H2122 cells treated with sotorasib for 2 or 24 hours. At each timepoint, cells were harvested and activation of the PI3K/AKT and MAPK pathways was assessed through western blot. (B) H2122 cells were treated with the stated dose of sotorasib, VVD-699 or a combination of both for 2 or 24 hours. At each timepoint, cells were harvested and activation of the PI3K/AKT and MAPK pathways was assessed through western blot. (C) H2122 cells treated with RMC-6236 for 2 or 24 hours. At each timepoint, cells were harvested and activation of the PI3K/AKT and MAPK pathways was assessed through western blot. (D) Anti-tumor efficacy of VVD-699, AMG-510, Binimetinib, or varying combination of both in H2122(KRAS^G12C^) xenografts. Data are shown as mean ± SEM; n=10 animals/group. Mice were dosed orally BID with 30 mg/kg VVD-699, QD with 30 mg/kg Binimetinib or QD with 100 mg/kg sotorasib. The same doses of each individual therapy were maintained when used in combination. (E) KPAR in vitro KPAR^G12C^ mouse cells were treated with 100 nM adagrasib, 500 nM VVD-699 or a combination of both for 2 or 24 hours. Cells were harvested and activation of the PI3K/AKT and MAPK pathways was assessed using western blot. (F) Survival of mice bearing KPAR^G12C^ orthotopic lung tumor treated with vehicle (n=6), 50 mg/kg QD adagrasib (n=5), 10 mg/kg QD VVD-844 (n=7) or the combination (n=5) for three weeks (grey area).

To confirm the relevance of these *in vitro* findings, we performed an *in vivo* efficacy study using H2122 xenografts, testing VVD-699 or the MEK inhibitor, binimetinib, in combination with sotorasib **(Fig. 5D)**. We find that VVD-699 provides a more profound benefit in combination with sotorasib than binimetinib, supporting our *in vitro* observations that sotorasib more effectively inhibits the MAPK pathway than the PI3K/AKT pathway. Additionally, while tumors treated with either doublet combination showed outgrowth, a triple combination showed strong and durable tumor regression. To evaluate the combination of RAS-p110α blocker with KRAS^G12C^ inhibitor in an orthotopic and syngeneic setting, we utilized the KPAR^G12C^ mouse lung cancer cell line (*25*), which shows a clear anti-growth effect with single agent VVD-844 in the subcutaneous setting (**Fig. S5D**). The in vitro behavior of the KRAS^G12C^ inhibitor, adagrasib, and VVD-699, alone and in combination was consistent with our observations in H2122 cells, suggesting that WT RAS dependent re-activation of the PI3K pathway is a common response to KRAS^G12C^ inactivation (**Fig. 5E**). Mice bearing KPAR^G12C^ orthotopically implanted lung tumors were treated with either adagrasib, the RAS-p110α blocker, VVD-844, or a combination for three weeks, followed by dosing release and close health monitoring until humane endpoint reached. VVD-844 alone had only modest impact on survival, while adagrasib treatment led to 40% of the mice surviving long term. However, the combination of VVD-844 and adagrasib was markedly more effective, with 100% of the mice showing complete and durable responses **(Fig. 5F)**.

## Discussion

The data presented here emphasize the importance of the interaction between RAS and the p110α RBD in the control of PI3K activity in cancer cells. Our chemoproteomic platform yielded multiple series of compounds that bind to cysteine 242 in the RBD of p110α, which lies adjacent to the binding interface with RAS. The residue is absent in other PI3K isoforms, ensuring isoform selectivity for p110α (*9, 26*). Amongst the C242 targeting compounds found in these studies, we identified not only inhibitors of the RAS/PI3K interaction, but also inducers of the interaction and neutral compounds that do not impact the interaction at all. This suggests a nuanced mechanism of small molecule action through binding this site, with the possibility of either inhibiting or promoting the binding depending on the precise nature of the compound bound to C242. This is reminiscent of previous work reporting that different mutations at a single residue in the RBD of p110α can either inhibit (K227A) or activate (K227E) PI3K activity (*6, 27*). While inhibitors and their possible role in cancer therapy are the main focus of the current work, inducers of the RAS/PI3K interaction could also have clinical utility for activating PI3K downstream of receptor tyrosine kinases, for example as a way of promoting tissue regeneration or treating insulin-resistant diabetes (*28, 29*).

Compounds such as VVD-699 block the interaction of RAS with p110α. In cells expressing an engineered G12V HRAS fusion protein where RAS can be acutely activated independently of other pathways, both VVD-699 and mutations in the RBD of p110α profoundly inhibit activation of PI3K. In RAS mutant cancer cells, this compound impairs PI3K activity, but to varying extents, likely reflecting the fact that direct, RAS-independent activation of PI3K by a variety of receptor tyrosine kinases also plays a very important role in PI3K regulation (*6*). Although these blockers by themselves have a minor impact on the proliferation of RAS mutant cancer cells in vitro, they do have greater inhibitory effects on the growth of RAS mutant or amplified tumors in vivo, both in xenografts and in orthotopic syngeneic mouse models, likely reflecting more stringent requirements for PI3K activity in these settings. The effectiveness of VVD-699 and similar compounds in vivo is markedly improved by combination with MEK inhibitors, as would be expected due to the need to block both MEK/ERK and PI3K effector pathways to achieve optimal control of tumor growth. In G12C KRAS mutant cells, our data supports that VVD-699 provides benefit in combination with KRAS-G12C inhibitors by blocking the rapid rebound in AKT phosphorylation, which is mediated, at least in part, via endogenous wild type HRAS and NRAS isoforms (*24*).

As anticipated from previous genetic experiments in mice (*16*), we found that mutant EGF receptor driven tumors showed sensitivity to compounds that disrupt the RAS/PI3K interaction. In addition, tumor cells over-expressing the related receptor tyrosine kinase HER2 showed a high degree of sensitivity to VVD-699, both in terms of PI3K activation and growth in vitro and in vivo. It is noteworthy that in HER2 over-expressing cancer cells, VVD-699 has a stronger impact on PI3K activity than does the pan RAS inhibitor RMC-6236, suggesting that there is a substantial RAS independent component to PI3K regulation downstream of HER2 that is nevertheless sensitive to compounds binding to the RBD of p110α. In support of a RAS independent mechanism of action for C242 ligands against HER2, we found that a C242 ligand that does not disrupt the RAS-PI3K interaction retained the ability to block HER2 signaling. This ligand also did not inhibit the interaction of p110α with RAS related isoforms RRAS, RAS2 (TC21) and MRAS, supporting that these isoforms are also not involved in HER2 signaling impacts of C242 liganding. Overall our data support an as yet unidentified, role for the p110α RAS Binding Domain in HER2 signaling, that can be perturbed by compounds binding to C242. Compared to RAS mutant tumors, the fact that HER2 over-expressing tumors are more consistently and profoundly sensitive to VVD-699 suggests that agents targeting the RBD of p110α could also find a role in their treatment.

Since the interaction of RAS with p110α appears to play a more important role in the regulation of PI3K by mutant RAS compared to its normal physiological regulation by growth factors (*6, 7*), it might be expected that normal regulation of PI3K would be less sensitive to C242 binding compounds. One of the leading toxicities seen with inhibitors of the kinase activity of PI3K in the clinic is impairment of insulin mediated control of blood glucose, which relies on PI3K activation via IRS1 with little input from RAS. The RBD targeting compounds reported here do not lead to hyperglycemia, unlike inhibitors of PI3K p110α kinase activity (*8*), suggesting a potential advantage of using compounds like VVD-699 over alpelisib in the treatment of RAS mutant, and potentially HER2 over-expressing, cancers. A compound related to VVD-699 will shortly be entering phase 1 trials in RAS mutant and other cancers, which should address whether the tolerability and efficacy of blocking the RAS Binding Domain of p110α seen in mice is translatable to the clinical setting.

## Acknowledgements

The authors thank Wuxi AppTec for compound synthesis and Crown Biosciences and Champions Oncology for animal study support. X-ray diffraction data were collected at the Advanced Light Source beamline 5.0.2. The Berkeley Center for Structural Biology is supported by the Howard Hughes Medical Institute, Participating Research Team members, and the National Institutes of Health, National Institute of General Medical Sciences, ALS-ENABLE grant P30 GM124169. The Advanced Light Source is a Department of Energy Office of Science User Facility under Contract No. DE-AC02-05CH11231.

## Author Contributions

N.R, J.D. and M.P.P conceptualized the project. J.E.K, N.R., T.M.K, G.M.S, D.S.W, J.P., I.M., J.D. and M.P.P supervised the project. Investigation and experiments were carried out by S.M.B, S.G., M.A.H., H.M., J.T., J.W., C.B., A.E.O., R.L., Y.L., M.P., H.P., I.M. S.R., A.N.S., E.J.W., T.E.W., E.A., K.B., J.C.B., J.M.C., K.H., B.D.H., M.I., K.N.L., W.L., J.M., M.K.P., J.J.S., M.T. and E.T. Writing, review and editing were undertaken by J.E.K., N.R., S.M.B., M.A.H., I.M., J.P., G.M.S., M.M.A., J.D and M.P.P.

---

**Supplemental Figure 1:**
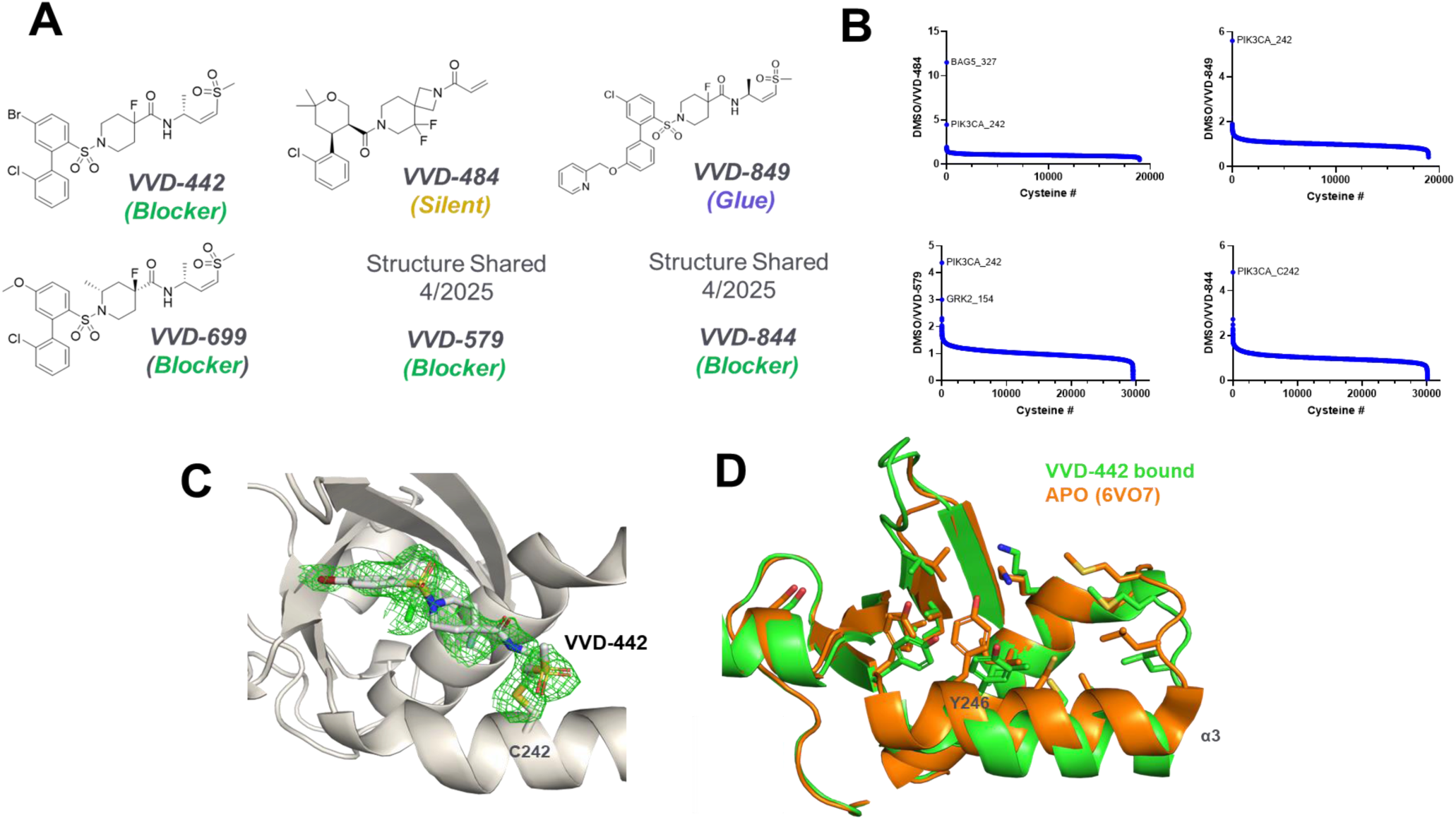
(A) Chemical structure of VVD ligands used in manuscript (B) Global proteomics selectivity of VVD-484, VVD-849, VVD-579, and VVD-844. Cells were treated with 10 µM of VVD-484, 2 µM of VVD-849 or VVD-579, or 1 µM of VVD-844, which is ∼25-fold over IC_50_s. 20,000-30,000 cysteine residues were measured using TMT quantification for each compound. (C) *F_O_-F_C_* omit map contoured at 2.5 σ showing continuous density for VVD-442 and a covalent bond formed with C242. (D) Superposition of the APO (PDB ID 6VO7) and liganded RAS-binding domain highlighting the shift in α3 and movement of Y246 required for compound binding.

**Supplemental Figure 2:**
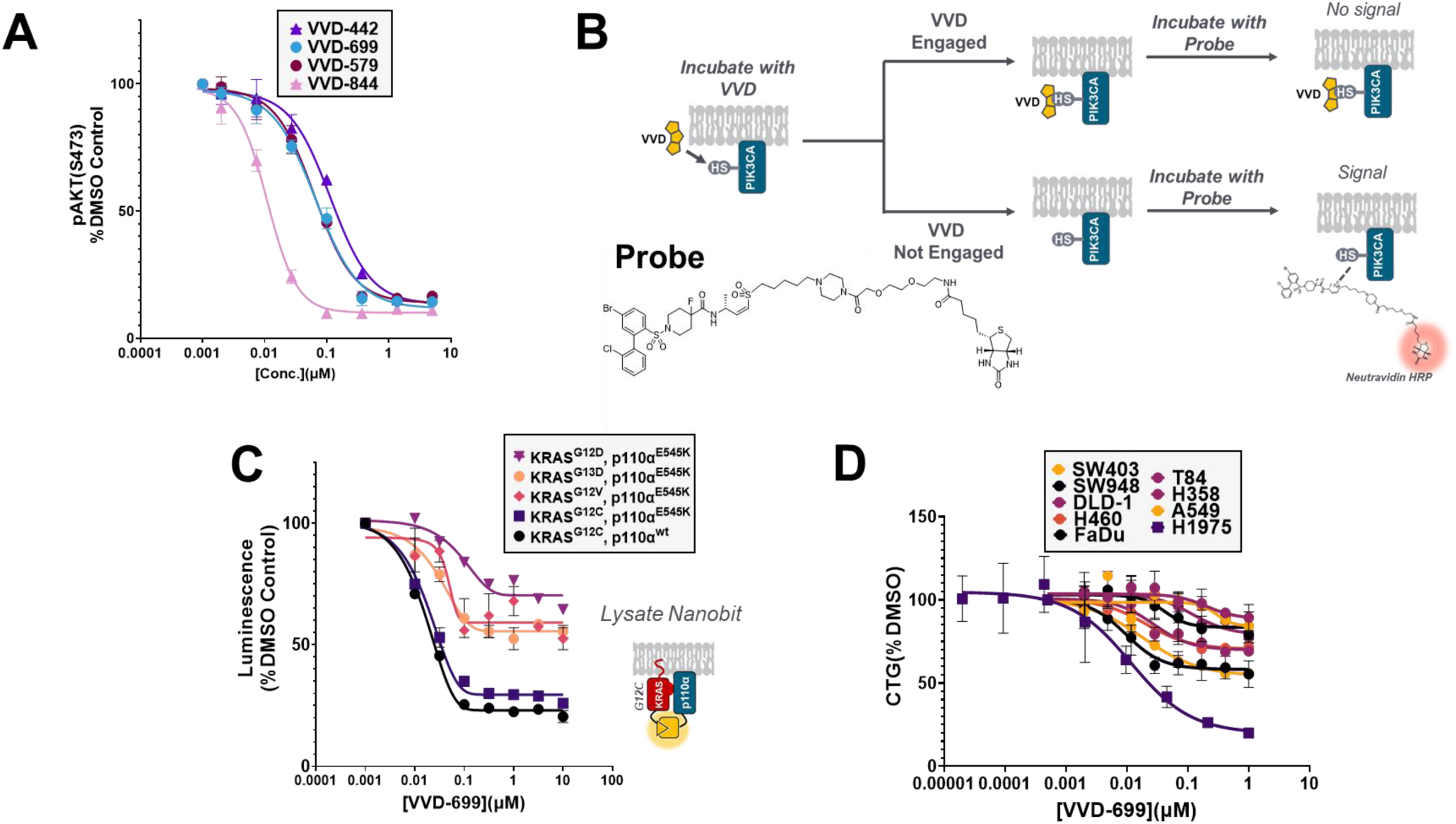
(A) H358 cells treated with a dose response of RAS-PI3K blockers for 30 minutes, followed by assessment of pAKT(S473) through HTRF (B) Diagram of pocket probe assay used to determine engagement of cysteine 242 on p110α (C) VVD-699 was screened for its ability to disrupt the interaction between p110α^E545K^ or p110α^WT^ and different KRAS mutants using the Nanobit protein-protein interaction assay in HEK293T cell lysates co-transfected with p85α and exposed to compound for 4 hours. (D) The ability of VVD-699 to inhibit growth of cells known to have high RAS activation was evaluated through 3D proliferation assays in which cells were treated with VVD-699 every 3 days for 9 days, followed by assessment for cell abundance using Cell Titer Glo.

**Supplemental Figure 3:**
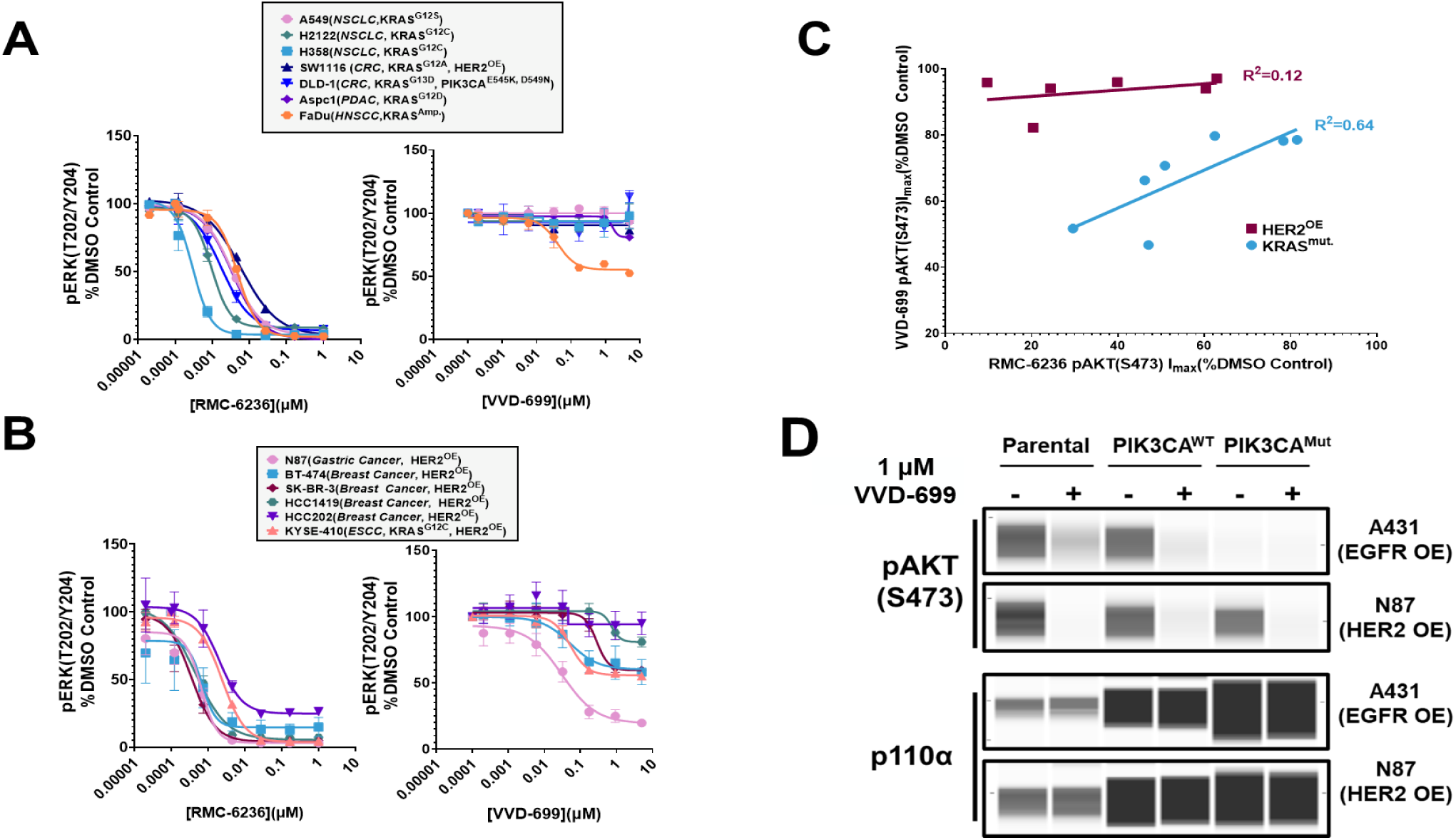
(A) KRAS hyperactive cell lines were treated with either RMC-6236 or VVD-699 for 2 hours, followed by assessment of pERK(T202/Y204) through HTRF. (B) HER2 overexpressing cell lines were treated with either RMC-6236 or VVD-699 for 2 hours, followed by assessment of pERK(T202/Y204) through HTRF. (C) Linear regression analysis of maximal inhibition seen with RMC-6236 or VVD-699 in KRAS^mut^(Fig. 3D) and HER^OE^(Fig. 3F) cells. (D) H1975(EGFR^mut^) or N87(HER2^OE^) cells were engineered to stably express p110α^RBDWT^ or p110α^RBDmut^ as well as the parental cell lines, were treated with VVD-699 for 2 hours. Samples were collected and pAKT(S473) and p110α levels were determined through Protein Simple western blotting.

**Supplemental Figure 4:**
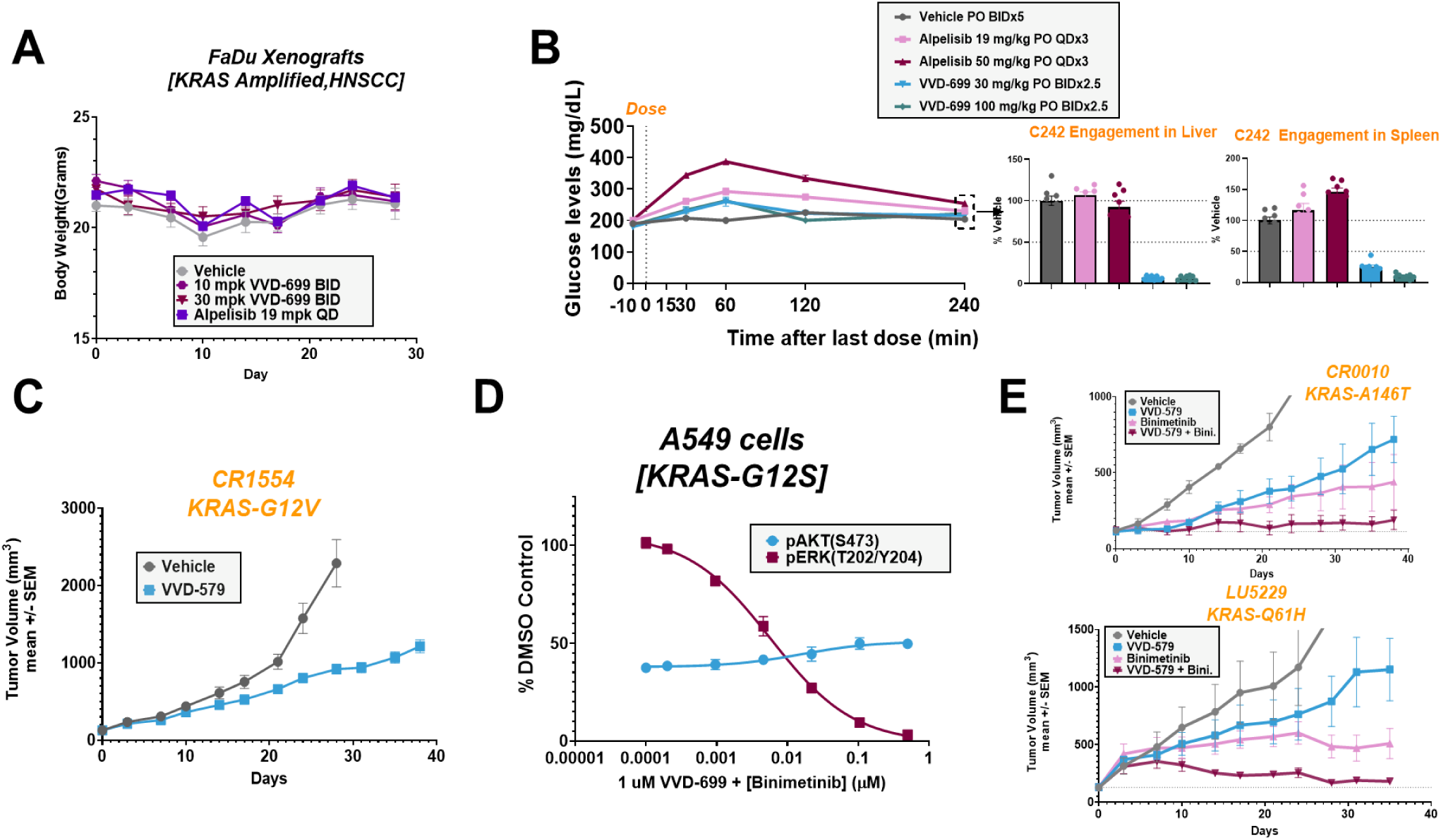
(A) Body weights of mice bearing FaDu(KRAS^amp^) xenografts that were treated with the indicated doses of VVD-699 or alpelisib. Data are shown as mean ± SEM; n=10 animals/group. (B) Mice were administered the indicated doses of alpelisib or VVD-699 for 3 days, followed by serial measurement of blood glucose levels for 4 hours post-last dose. At the conclusion of the time-course, spleen and liver were collected from the animals and p110α-C242 engagement was measured through mass spectrometry. (C) Anti-tumor efficacy of VVD-579 in KRAS^G12V^ PDX model CR1554. Data are shown as mean ± SEM; n=3 animals/group. Mice were dosed orally BID with 30 mg/kg VVD-579. (D) A549(KRAS^G12S^) cells treated with a static 1 µM dose of VVD-699 and a dose response of binimetinib for 2 hours. Samples were collected and pAKT(S473) and pERK(T202/Y204) levels were determined through ProteinSimple western blotting. (E) Anti-tumor efficacy of VVD-579, Binimetinib, or a combination of both in CR0010(KRAS^A146T^) (Top) or LU5229(KRAS^Q61H^) (Bottom) PDX models. Data are shown as mean ± SEM; n=3 animals/group. Mice were dosed orally BID with 30 mg/kg VVD-579, QD with 30 mg/kg Binimetinib or a combination of both.

**Supplemental Figure 5:**
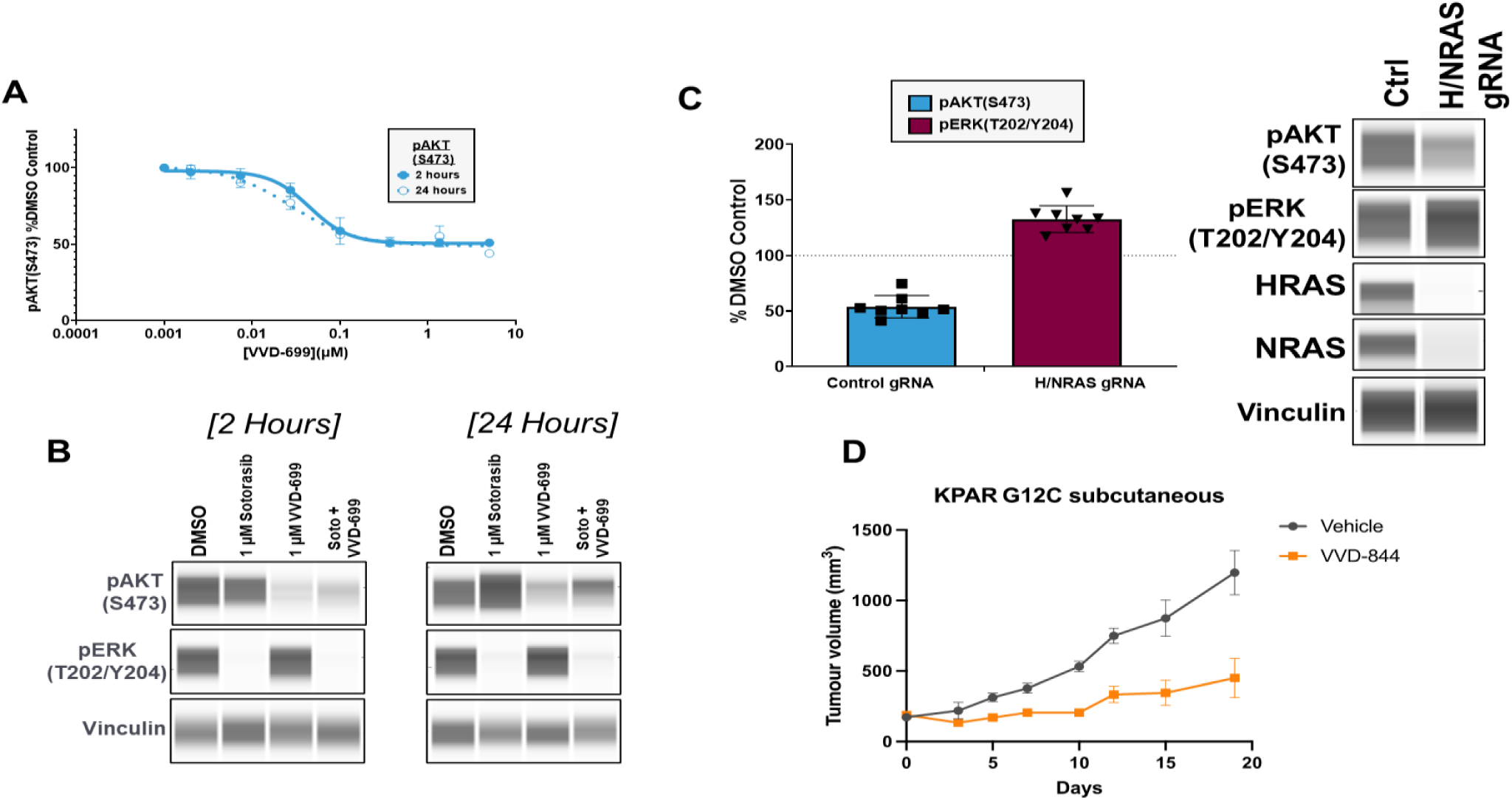
(A) H2122(KRAS^G12C^) cells were treated with a dose response of VVD-699 for 2 or 24 hours. At each timepoint, samples were collected and pAKT(S473) was measured through HTRF. (B) H2122(KRAS^G12C^) cells were treated with indicated compounds for 2(B) or 24(C) hours. Samples were collected and pAKT(S473) and pERK(T202/Y204) levels were compared against through ProteinSimple western blotting. (C) H2122(KRAS^G12C^) cells were transfected with CRISPR control guides or guides targeting HRAS and NRAS for 72-96 hours. Samples were collected and pAKT(S473) and pERK(T202/Y204) levels were compared against through ProteinSimple western blotting. Knockdown of HRAS and NRAS was also confirmed through ProteinSimple western blotting. (D) Anti-tumor efficacy of VVD-844 in subcutaneous KPAR^G12C^ tumors. Data are shown as mean ± SEM; n=8-9 mice/group. Mice were dosed orally QD with 10 mg/kg VVD-844.

**Supplemental Table 1:** Summary of VVD ligands described in this manuscript. K_i_ and k_inact_ values for VVD-699 determined with recombinant p110α using intact protein MS. All additional values determined using methodology described elsewhere in this manuscript. H358 cells were treated with compound for 30 minutes prior to collection and evaluation of pAKT(S473) via HTRF.

| | $k_{inact}$ | $K_i$ | $k_{inact}/K_i$ | Mass Spec. | Pocket Probe | KRAS<sup>G12C</sup>:p110α Lysate Nanobit | | pAKT(S473) [H358 cells] | |
| --- | --- | --- | --- | --- | --- | --- | --- | --- | --- |
| ID | s<sup>-1</sup> | μM | M<sup>-1</sup> s<sup>-1</sup> | TE<sub>50</sub> (μM) | TE<sub>50</sub> (μM) | IC<sub>50</sub> (μM) | I<sub>max</sub> (%DMSO) | IC<sub>50</sub> (μM) | I<sub>max</sub> (%DMSO) |
| VVD-442 | ND | ND | ND | 0.089 | 0.11 | 0.11<sup>a</sup> | 76%<sup>a</sup> | 0.115 | 85% |
| VVD-699 | 0.018 | 40.7 | 445 | 0.05 | 0.075 | 0.036<sup>a</sup> | 78%<sup>a</sup> | 0.066 | 86% |
| VVD-579 | ND | ND | ND | ND | 0.06 | 0.03<sup>a</sup> | 75%<sup>a</sup> | 0.063 | 84% |
| VVD-844 | ND | ND | ND | ND | 0.009 | 0.004<sup>a</sup> | 86%<sup>a</sup> | 0.011 | 90% |
| VVD-484 | ND | ND | ND | 0.44 | 0.50 | 0.59<sup>b</sup> | -77%<sup>b</sup> | ND | ND |
| VVD-849 | ND | ND | ND | 0.063 | 0.1 | 0.02<sup>a</sup> | -378%<sup>a</sup> | 0.04 | -379% |

<sup>a</sup>4 hour compound incubation, p85α co-transfected
<sup>b</sup>1 hour compound incubation

**Supplemental Table 2.**
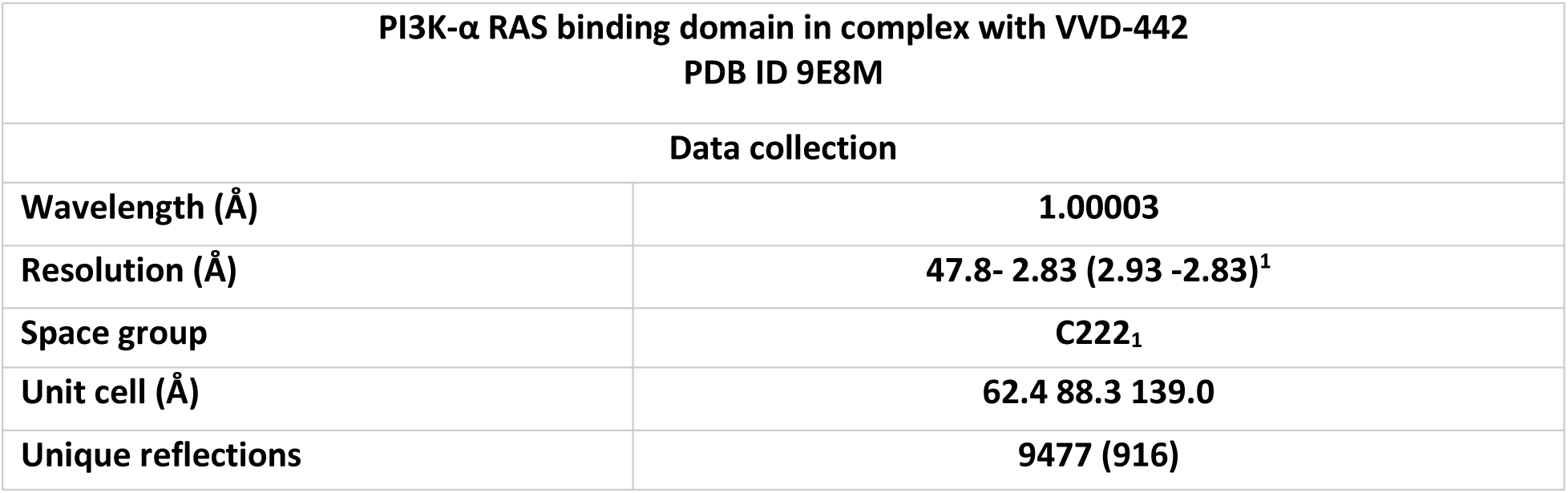
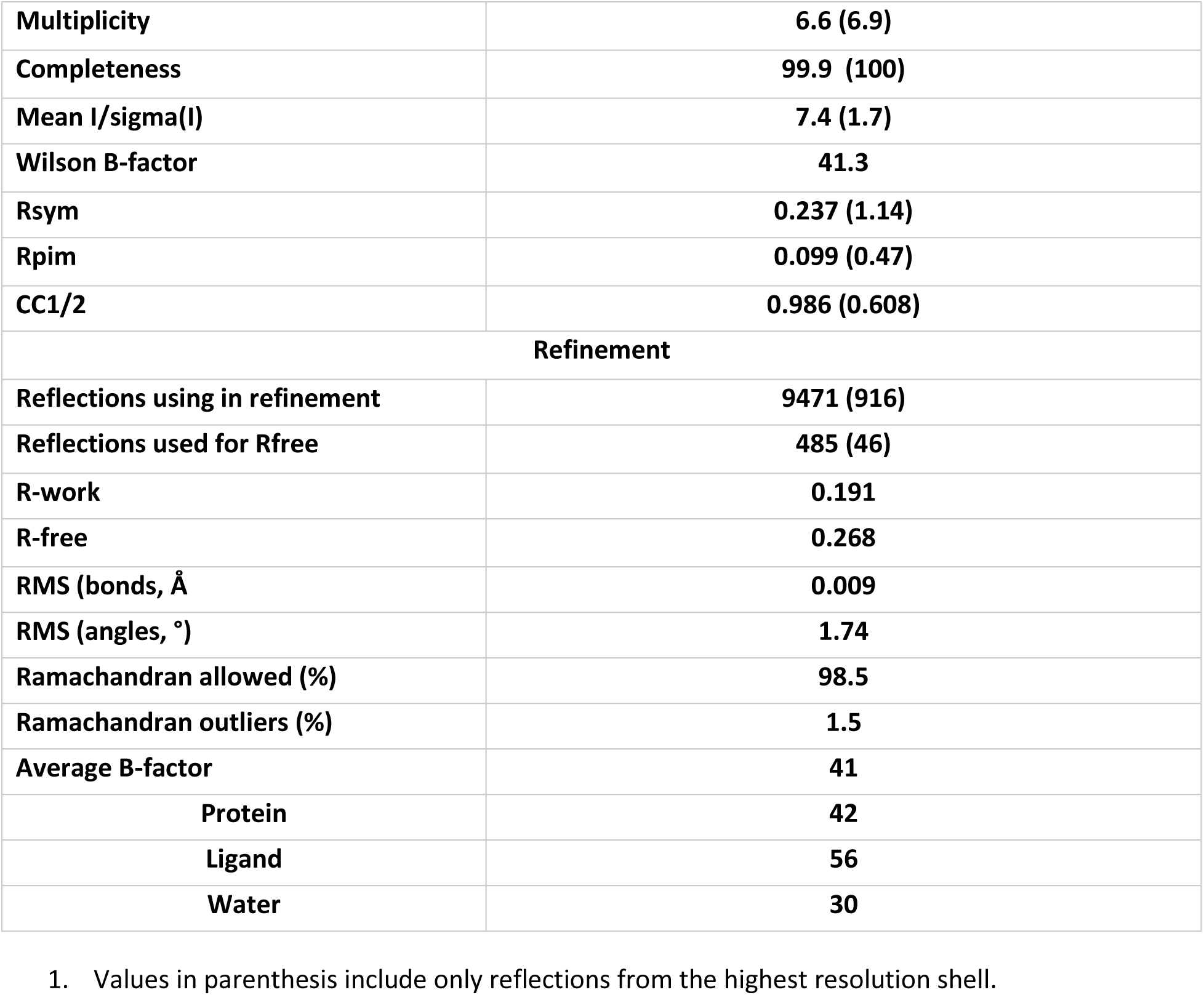

| PI3K-α RAS binding domain in complex with VVD-442<br>PDB ID 9E8M | |
| --- | --- |
| Data collection |  |
| Wavelength (Å) | 1.00003 |
| Resolution (Å) | 47.8- 2.83 (2.93 -2.83)<sup>1</sup> |
| Space group | C222<sub>1</sub> |
| Unit cell (Å) | 62.4 88.3 139.0 |
| Unique reflections | 9477 (916) |
| **Multiplicity** | **6.6 (6.9)** |
| **Completeness** | **99.9 (100)** |
| **Mean I/sigma(I)** | **7.4 (1.7)** |
| **Wilson B-factor** | **41.3** |
| **Rsym** | **0.237 (1.14)** |
| **Rpim** | **0.099 (0.47)** |
| **CC1/2** | **0.986 (0.608)** |
| **Refinement** |  |
| **Reflections using in refinement** | **9471 (916)** |
| **Reflections used for Rfree** | **485 (46)** |
| **R-work** | **0.191** |
| **R-free** | **0.268** |
| **RMS (bonds, Å)** | **0.009** |
| **RMS (angles, °)** | **1.74** |
| **Ramachandran allowed (%)** | **98.5** |
| **Ramachandran outliers (%)** | **1.5** |
| **Average B-factor** | **41** |
| **Protein** | **42** |
| **Ligand** | **56** |
| **Water** | **30** |

1. Values in parenthesis include only reflections from the highest resolution shell.

## Materials and Methods

### Cell Lines and culture

#### Cell line source

Jurkat (ATCC; catalogue # TIB-152)

HEK293T (ATCC; catalogue # CRL-3216)

H358 (ATCC, catalogue # CRL-5807)

A549 (ATCC, catalogue # CCL-185)

AsPC-1(ATCC, catalogue # CRL-1682)

T24(ATCC, catalogue # HTB-4)

SW1116(ATCC, catalogue #CCL-233)

H1975(ATCC, catalogue # CRL-5908)

H2122(ATCC, catalogue # CRL-5985)

T84(ATCC, catalogue # CCL-248)

SW403(ATCC, catalogue # CCL-230)

DLD-1(ATCC, catalogue # CCL-221)

FaDu(ATCC, catalogue # HTB-43)

HCC-202(ATCC, catalogue # CRL-2316)

BT-474(ATCC, catalogue # HTB-20)

N87(ATCC, catalogue # CRL-5822)

SK-BR-3(ATCC, catalogue # HTB-30)

HCC1419(ATCC, catalogue # CRL-2326)

KYSE-410( Sigma Aldrich, catalogue #94072023-VL)

KPAR1.3 KRAS^G12C^ cells (herein KPAR^G12C^) were generated as previously described ( DOI: 10.1158/0008-5472.CAN-22-0325)(*30*)

MEFs p110α RBD wt HRAS^G12V^-ER and MEFs p110α RBD mut HRAS^G12V^-ER were generated as previously described (DOI: 10.1016/j.cell.2007.03.051)(*6*)

#### Cell Culture conditions

Cell lines were maintained according to the vendor handling instructions

### Compound treatment

Cells were dosed with the indicated dose response or set dose of compound using an HP Digital Dispenser.

For in vivo studies, the following vehicles were used for the respective compounds:

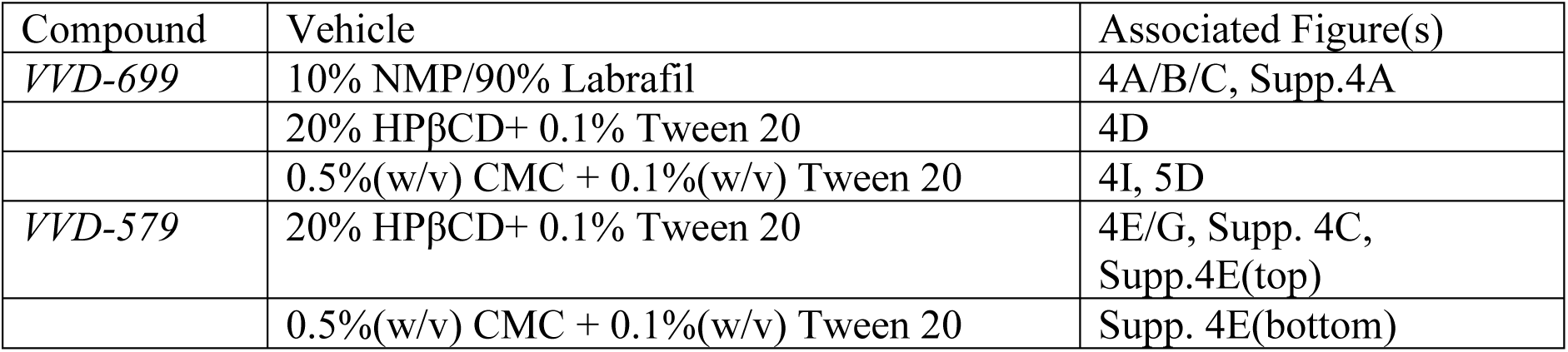
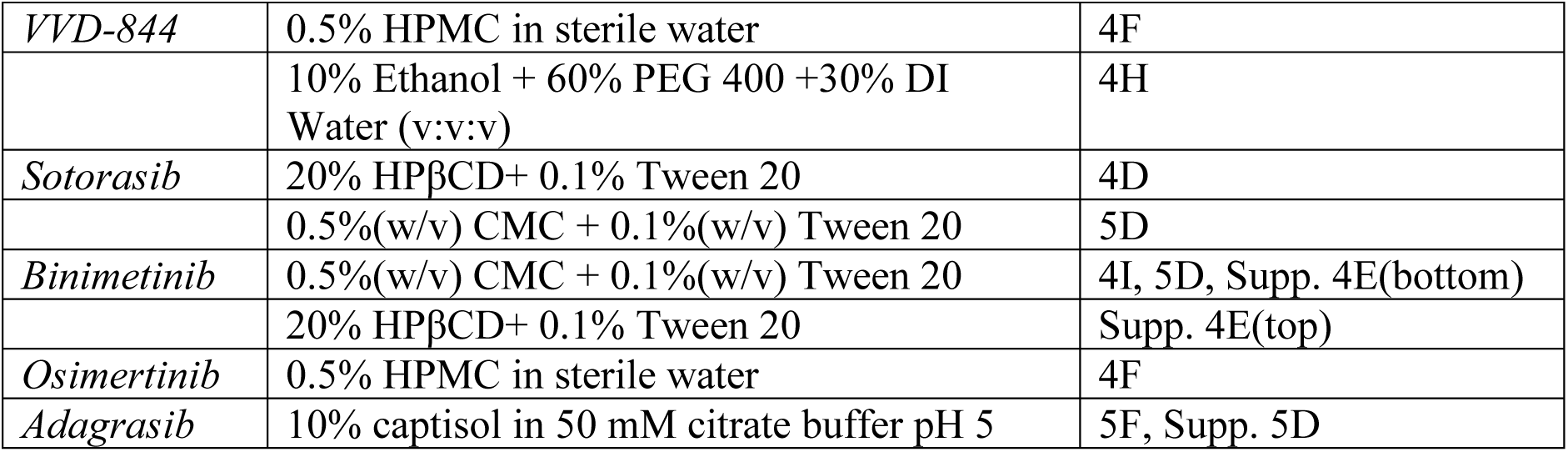

### Targeted Chemoproteomics

Chemoproteomics samples were prepared as previously described and detailed briefly below(*11*).

#### In Vitro target engagement

Jurkat or Karpas-299 cell pellets were lysed by probe sonication in Dulbecco’s phosphate buffered saline (D-PBS) and, following protein concentration normalization, 500 µg of total protein per well was aliquoted into a 96-well plate. Lysates were incubated with compound for 1 h followed by iodoacetamide-desthiobiotin (IA-DTB) probe-labeling of all solvent exposed cysteines. Protein was acetone precipitated then resuspended in freshly prepared 9M urea with 50 mM ammonium bicarbonate. Proteins were reduced and alkylated by the addition of DTT and iodoacetamide, buffer exchanged with Zeba de-salting plates, then digested with trypsin for 1h. IA-DTB labeled peptides were isolated with streptavidin agarose resin and enriched cysteine-containing peptides were eluted by the addition of 50% acetonitrile (ACN) with 0.1% Formic Acid (FA) and dried until ready for analysis. For VVD-849, probe labeled samples were TMT labeled (detailed below) and target engagement quantified by targeted TMT, as previously described. (*32*)

#### In Vivo target engagement

Snap-frozen tissues in bead-beater tubes were placed on ice and homogenized in cold Pierce RIPA lysis buffer with 30 s of bead beating at 4 °C. Samples were sonicated and cleared by centrifugation, protein was normalized, and 300 µg per sample aliquoted into a 96-well plate. Samples were prepared as described above, starting with IA-DTB probe-labeling.

#### Liquid chromatography – tandem mass spectrometry analysis

Probe-labeled peptides were concentrated onto an Acclaim PepMap100 C18 loading column (Thermo, DX164564, 100 µm x 2 cm, 5 µm particle size) and separated on a custom made C18 nanoviper analytical column (Thermo, 75 µm x 15 cm, 2 µm particle size) using a Dionex Ultimate 3000 nano-LC (Thermo). For in vitro experiments, peptides were separated using an 8.7 min gradient going from 6 to 32.5% B solvent (96.4% ACN, 3.5% dimethylsulfoxide (Pierce, PN 20688), 0.1% FA) mixed with A solvent (96.4% water, 3.5% dimethylsulfoxide, 0.1% FA). In vivo samples were separated using the same gradient, but over 12.7 min.

Peptides were analyzed by parallel reaction monitoring (PRM) mass spectrometry using product ion scan mode on the Thermo Exploris 120 orbitrap mass spectrometer in positive ion mode with a spray voltage of 2500V and 375 °C ion transfer tube temperature. Precursor ions were fragmented and measured using a normalized collision energy of 25%, 0.7 m/z Q1 resolution, 30,000 orbitrap resolution, and 70% RF lens. In vitro datasets were acquired with a 200% normalized AGC for all peptides and in vivo datasets were acquired with custom normalized AGC up to 3000% for PIK3CA peptides. Data was acquired using a scheduled method with 1.2 min windows and dynamic injection time mode with a minimum of seven points across the peak.

#### Parallel Reaction Monitoring (PRM) data analysis

Target engagement (PIK3CA_C242) was measured by monitoring peptide LCVLEYQGK (688.8839 m/z) and additional PIK3CA peptides were monitored to ensure target engagement rather than a protein level change. Retention time standard (RTS) peptides were also included in the method and were used for global normalization for each sample. Mass Spectrometry data was analyzed using Skyline v.22.2.0.255 (MacCoss Lab, University of Washington). Peptide quantification was performed by summing the peak areas corresponding to four to six fragment ions. Fragment ions were pre-selected from an in-house generated reference spectral library. RTS peptides were used to normalize for sample variability. Percent target engagement for each peptide was determined by comparing the average peptide AUC of the compound treated group to the average peptide AUC of the control (DMSO or vehicle treated) group.

### Global Proteomic Selectivity Profiling

#### Live cell target engagement

Five million jurkat cells (in RPMI with 10% FBS) per well were arrayed into a 96-well plate and treated with compound for 2 h. Following compound wash out, cells were lysed by sonication in phosphate buffered saline (PBS) and incubated with IA-DTB for one hour at room temperature. Cysteine peptide enrichment was performed as described above. The resulting dried peptides were resuspended in 0.2 M EPPS pH 8.5 in 30% anhydrous ACN and labeled with 55 µg TMTpro 18plex reagent (Thermo, A52045). Samples were quenched with hydroxylamine. The combined samples were desalted on a Biotage evolute express ABN plate and separated by high pH reversed phase fractionation. 96 fractions were recombined to 12 fractions and analyzed by LC-MS/MS. Three 18-plexes were used to generate this data and compounds were paired as follows: VVD-699/VVD-844, VVD-442/VVD-579, and VVD-484/VVD-849.

#### Liquid chromatography – tandem mass spectrometry analysis

Cysteine enriched peptides were loaded onto an Acclaim PepMap100 C18 loading column (listed above) and separated on an Acclaim PepMap C18 analytical column (Thermo, 75 µm x 25 cm, 2 µm particle size) using a Dionex Ultimate 3000 nano-LC (Thermo). Separation was using an 83.5 min gradient from 6 to 30% solvent B for VVD-442, VVD-699, VVD-844, and VVD-579 and a 171 min gradient from 6 to 30% solvent B for VVD-484 and VVD-849. Global TMT data was acquired on an Orbitrap Fusion Lumos Tribrid MS (Thermo) using the SPS MS3 workflow.

#### Global TMT data analysis

Data was searched with MassPike software package (GFY development team, GFY Core Version 3.4) using the Human FASTA database with a precursor mass tolerance of 30 ppm and a fragment ion tolerance of 0.3 m/z and filtered using a 2% false discovery rate. Peptide spectral match results with an XCorr > 1 and a summed DMSO TMT signal > 20 were converted to cysteine site level data, then filtered to remove sites with >40% CV in the DMSO control samples. Each compound was prepared with a dose response above the selected concentration (∼25-fold over average IC_50_s) and sites that did not exhibit a dose response were filtered from the dataset (∼300 out of 30,000 sites filtered from VVD-442, VVD-699, VVD-844, and VVD-579 datasets and ∼30 out of 20,000 sites filtered from the VVD-484 and VVD-849 dataset). Global data normalization was applied and ratio of average DMSO AUC to compound treated AUC calculated to generate selectivity data plots.

### Pocket Probe

#### PIK3CA Pocket Probe Generation

A covalent ligand that binds potently and selectively to cysteine-242 of PIK3CA was identified. An exit vector from this covalent ligand was identified and linked to D-Biotin to generate a PIK3CA biotin probe that was used under the assay conditions. This probe has a proteomic TE_50_ of 100 nM in Jurkat cell lysate.

#### Pocket probe PIK3CA target engagement in Jurkat and H358 cell lysates

Cell pellets were thawed and diluted in cold Dulbecco’s phosphate buffered saline to a concentration of 300 × 10^6^/12 mL(Jurkat) or 300 × 10^6^/20 mL(H358). Cells were lysed via sonication, and lysate was plated (50 µL/well) in a 96-well plate. The cell lysates were then treated with compound using the HP Digital Dispenser and allowed to incubate at room temperature for 1 hour(Jurkat) or 2 hours(H358). Next, the biotin-conjugated pocket probe was added to each well and allowed to incubate for an additional hour at room temperature. 75 uL of dilution buffer was added to each well, followed by centrifugation and storage at -80°C. On the day of the assay, the lysate was thawed and added to a 96-well plate pre-coated with capture antibody. Following a 1.5 hour incubation, the plate was washed with 100 µL/well PBST, and 20 µL/well of NeutrAvidin-HRPwas added and incubated for 1 hour at room temperature. The plate was then washed 4 times with PBST, 100 µL/well. HRP-substrate, 20 µL/well, was added to the plate prior to the CLARIOstar Plus luminescence read.

#### Analysis

To account for non-specific assay signal, first a qualitative assessment was performed to assess stability of the bottom of the curve, i.e. no additional loss in signal at the three highest doses. If so, this residual signal was deemed to be the floor of the assay and the signal at the highest dose of compound, 50 µM, was subtracted from all other datapoints. The treatment samples were then normalized to the DMSO treated samples and the TE_50_ was determined using GraphPad Prism software (part of Dotmatics, Boston MA).

### RAS-p110α Nanobit

#### HEK293T Lysate format

HEK293T cells plated in full media in a T225 flask were transfected with pBiT2.1-N [TK/SmBiT]_KRAS^mut^, pBiT1.1-N [TK/gBiT]_PI3KCA^wt^/^E545K^ using Lipofectamine 3000. HSV-TK-p85α was also co-transfected where indicated.

The cells then incubated for 48 hours at 37^°^C with 5% CO_2_. The cells were then harvested and lysed through probe sonication, followed by storage of the cell lysate at -80^°^C.

On the day of the assay, the lysate was thawed and 20 µL/well of lysate was distributed to a 384 well plate, as well as 20 uL of untransfected HEK293T cell lysates, which served as a negative control. The plate was then treated with compound using the HP Digital Dispenser and following a 1-hour(p85α not co-transfected) or 4-hour(p85α co-transfected) incubation at RT on an orbital shaker, 4 µL of Nano-Glo reagent was added to each well and luminescence was read using the CLARIOstar Plus (BMG LABTECH, North Carolina).

#### H358 Live Cell Format

To generate the pBiT2.1-CMV-Smbit-RAS expression plasmids, full length cDNA for amplified for the respective isoforms was using PCR and cloned in-frame into the pBiT2.1-N [TK/SmBiT] vector. To generate the pBiT1.1-CMV-Lgbit-PIK3CA WT expression plasmid, PIK3CA was amplified using PCR and cloned in-frame into the pBiT1.1-N [TK/LgBiT] vector. For both expression plasmids, Gibson cloning was utilized to replace the TK promoter with the CMV promoter. To generate pBiT1.1-CMV-Lgbit-PIK3CA RBD, in which PIK3CA can no longer interact with RAS, two point mutations on the RAS binding domain(RBD) of PIK3CA (T208D and K227A) were introduced using the Q5 site-directed mutagenesis kit(*6*).

20,000 H358 cells/well were plated in full media in a 96 well plate and incubated overnight at 37^°^C with 5% CO_2_. The following morning, the cells were transfected with pBiT2.1-CMV-Smbit-KRAS G12C(50ng/well), pCMV6-p85β (100ng/well) and pBiT1.1-CMV-Lgbit-PIK3CA WT or pBiT1.1-CMV-Lgbit-PIK3CA RBD(100ng/well) using Lipofectamine 3000. The transfected cells then incubated for 24 hours at 37 °C with 5% CO_2._ Following this incubation, cell media was removed and replaced with Opti-MEM media, followed by compound treatment using the HP Digital Dispenser. The plate then incubated 37 °C with 5% CO_2_ for 2 hours. Following this incubation, Nano-Glo Live substrate was diluted 1:20 in Nano-Glo LCS Buffer, pre-warmed to 37 °C. 25 µl of the mixture was added to each well and gently mixed by hand. The plate was then incubated for 20 minutes at 37°C and then luminescence was determined us the Clariostar Plus plate reader (BMG LABTECH, North Carolina, USA).

### pAKT(S473) and pERK1/2(T202/Y204) HTRF assays

Cells were harvested, centrifuged at 1000 rpm for 5 minutes and resuspended in media containing 1% FBS. 100 µL of cell suspension was plated into each well of a 96 well plate, corresponding to 25000-40000 cells/well, optimized on a per cell line basis. The plate was incubated overnight at 37 °C with 5% CO_2_ to allow the cells to adhere to the plate. The following day, the cells were treated with a dose response of compound. The plate was incubated for the indicated amount of time at 37 °C with 5% CO_2_. Following this incubation, media was removed and HTRF lysis buffer, was added to each well, followed by an incubation for 45-60 minutes at room temperature. The lysate was then transferred to an HTRF plate pre-loaded with HTRF antibody mix. Additionally, wells were loaded with lysis buffer only and HTRF antibody mix as a negative control. The plate was then sealed and allowed to incubate overnight at room temperature. The following morning, fluorescence intensity was measured on the Clariostar Plus plate reader Company, (BMG LABTECH, North Carolina, USA).

The ratio of acceptor/donor fluorescence of all wells was calculated using the following formula:

<u>Fluorescence Ratio</u>=Signal 665 nM/Signal 620 nM

The average ratio of the negative control wells was calculated and subtracted from all wells to account for background signal. Then the fluorescence ratio from each compound treated well was normalized to the fluorescence ratio of the DMSO treated well (100% activity)

### ProteinSimple Western Blot Analysis

#### In vitro

##### Cell Treatment

Cells were harvested, centrifuged at 1000 rpm for 5 minutes and resuspended in media with 1% FBS. 100 µL of cell suspension was plated into each well of a 96 well plate, corresponding to 30,00 or 40,000 cells per well, depending on the cell line. The plate was incubated overnight at 37 °C with 5% CO_2_ to allow the cells to adhere to the plate. The following day, the cells were treated with a dose response, or set dose, of compound using the HP Digital Dispenser for the indicated amount of time. For the 24-hour timepoint, the plate was incubated for 22 hours at 37 °C with 5% CO_2_. The media was then removed from the plate and fresh media was added, followed by a second treatment of compound, using the previously described dosing format, and an additional 2 hour incubation at 37 °C with 5% CO_2_.

##### Cell Harvesting and lysate preparation

Following the incubation period, media was removed and cells were lysed with 16 µL of supplemented RIPA buffer. Immediately after harvesting, 8 µL of sample was moved to a PCR strip tube pre-loaded with 2 µL of ProteinSimple 5x Master Mix and the mixture was incubated in the thermocycler for 10 minutes at 95°C. The remaining lysate was used to determine protein concentration through BCA assay according to manufacturer’s instructions (Fisher Scientific, Cat # PI23225). After boiling, using the determined protein concentration, lysates were normalized using 1X sample buffer(5x diluted with prepared RIPA lysis buffer).

#### In vivo

Tumor samples were collected in bead beater tubes and snap frozen at the time of collection. Samples were kept on ice and each tube was filled 350 µL of lysis buffer(RIPA buffer, Benzonase, MgSO_4_, HALT and Okadaic Acid), followed by homogenization by bead beating at 4 °C for 30 seconds. Following a hard spin to clear insoluble material, 50 µL of sample from each tube was transferred to a 96 well plate and kept on ice. 5 µL of each sample was then transferred to a new well, pre-loaded with 20 µL of lysis buffer and mixed. A BCA assay was then performed on the samples to determine protein concentration. Following normalization with lysis buffer, 8 µL of homogenate was transferred to PCR tubes pre-loaded with 2 µL of 5x sample buffer. The tubes were then briefly mixed and boiled at 95°C for 10 minutes and kept on ice.

#### Analysis using Jess automated western blotting (in vitro and in vivo)

The lysates were probed with CST #4060 (pAKT(S473)), CST #4370 (pERK1/2(T202/Y204)), CST #4650 (Vinculin)(Loading Control)

In all cases, manufacturer provided Rabbit-HRP secondary antibody, pre-diluted by the manufacturer, was used following incubation with primary antibody.

Each capillary was analyzed on an individual basis by considering the graph of Chemiluminescence relative to expected MW of the proteins being detected. Each protein being detected was then normalized to the Vinculin signal, serving as a loading control, within the capillary using the follow formula:

<u>Normalized Signal</u>: AUC of Protein of Interest(pAKT(473) or pERK1/2(T202/Y204)/ AUC of Vinculin

Then the normalized signal of each protein of interest from each treated well was normalized to the matching normalized signal from the DMSO treated well (100% activity).

### Traditional Western Blot Analysis

20 μg of cell lysate was subjected to electrophoresis in 4-12% NuPAGE Bis-Tris gels (Life Technologies) followed by transfer to nitrocellulose membrane. Lysates were probed with phospho-ERK (T202/Y204, #9101), ERK (#9107), phospho-AKT (S473, #9271) and AKT (#2920) from CST and Vinculin (V4505) from Sigma. Bound primary antibodies were incubated with secondary antibodies compatible with infrared detection at 700 nm and 800 nm. Membranes were scanned using the Odyssey Infrared Imaging System (Odyssey, LICOR).

### 3D#viability assays

Cells were harvested, centrifuged at 1000 rpm for 5 minutes, and resuspended in media with optimized FBS content to promote cell growth. The following conditions were used for the identified cell lines:

*10% FBS*: FaDu, N87, SK-BR-3, HCC1419, HCC202, KYSE-410, BT-474, SW403, SW948, H358

*5% FBS*: DLD-1, A549

*1% FBS*: H460, T84

100 µL of cell suspension was plated into a 96 well low attachment plate and incubated overnight at 37°C to allow the cells to form a sphere. The cells were then treated with a dose-response curve of VVD-132699 using the HP Digital Dispenser. The plate was then returned to the 37°C incubator. Every 3 days, the compound was refreshed by treating cells with the same dose-response curve of VVD-132699, using the HP Digital Dispenser and then returned to the 37°C incubator. For all cell lines except N87, which was harvested on Day 6, 100 µL of 3D CellTiter-Glo (CTG) was added to each well, mixed vigorously for 5 minutes, and then incubated at room temperature for 25 minutes on Day 9. Fluorescence intensity was then measured on the CLARIOstar plate reader (BMG LABTECH, North Carolina, USA).

### A549 p110α-C242S clone generation

Introduction of the C242S mutation into A549 cells was achieved through CRISPR/Cas9 technology. Briefly, A549 cells were transfected with RNP complexes, containing Cas9 and guide RNA, and ssDNA using the Lonza 4D Nucleofector X unit. After electroporation, cells were moved to a 6 well plate and incubated for 48-72 hours at 37 °C with 5% CO_2_. To confirm the C242S mutation, cells from one well were harvested and subjected to DNA sequencing. The desired mutation was confirmed at a low frequency, therefore, single cell clones were generated. This was achieved by seeding single cells in a 384 well plate. As the clones grew, they were progressively expanded until enough cells were present to seed a 24 well plate. At this point, cells were again subjected to DNA sequencing, at which point multiple clones carrying homozygous C242S were identified. Following this confirmation, cells were expanded and banked.

### In vitro kinase assay

The assay was performed using ADP-Glo Kinase assay reagents (Promega). It measures kinase activity by quantitating the ADP amount produced from the enzymatic reaction. The luminescent signal from the assay is correlated with the amount of ADP present and is directly correlated with the amount of kinase activity. The compounds were diluted in 2.5 % DMSO and 5 µl of the dilution was added to a 25 µl reaction so that the final concentration of DMSO is 0.5 % in all of reactions. The enzymatic reactions were conducted at 30 ° C for 45 minutes. The 25 µl reaction mixture of PI3 kinase contains 40 mM Tris, pH 7.4, 20 mM MgCl_2_, 0.1 mg/ml BSA, 2.5 µM ATP, kinase substrate and the enzyme. After the enzymatic reaction, 25 µl of ADP-Glo reagent was added and incubated for 45 min at room temperature followed by another 30 min incubation with 50 µl of kinase detection mixture. Luminescence signal was measured using a BioTek *Synergy 2* microplate reader.

### Protein expression and purification

The DNA sequence for RBD domain for crystallization (residues 157-300 was synthesized and cloned into pET28b with an N-terminal hexa-histidine SUMO tag. The gene for PI3K residues 105-1048 was synthesized and cloned into pFastBac1 with an N-terminal 6xHis tag followed by a TEV cleavage site. For *E. coli* expression, plasmids were transformed into BL21 DE3 Star. Large scale cultures were grown in 2XYT medium at 37C to an OD600 of 0.6 and induced with 0.4 mM IPTG at 18C overnight. Full lengthPI3KCa was expressed in ExpiSf9 cells using baculovirus mediated expression in the EmBacY viral genome. RBD domain for crystallization was purified using the following procedure. Cells were resuspended in 50 mM HEPES pH 8, 250 mM NaCl, 10% glycerol, 10 mM imidazole and 1mM TCEP and lysed using a Microfluidizer. The lysate was cleared by centrifugation at 25,000 x g for 45 minutes before being loaded onto 5 ml of pre-equilibrated NiNTA resin. The resin was washed with 500 ml lysis buffer and eluted with lysis buffer supplemented with 250 mM imidazole. The SUMO tag was removed with SUMO protease and the sample was dialyzed against lysis buffer. Uncleaved protein was removed with an additional NiNTA purification before additional purification with a Superdex S200 column equilibrated in 25mM HEPES, pH 7.5, 50mM NaCl 3 mM TCEP. Full length PIK3CA was purified in a similar manner, omitting the proteolytic cleavage steps.

### Crystallization and structure determination

Purified PI3K RBD was incubated with VVD-442 in 25 mM Tris pH 7.5, 150 mM NaCl, 5% glycerol, 1 mM TCEP, 2% DMSO and the reaction was monitored by intact protein MS. Upon completion, the protein sample was buffer exchanged into 25 mM Tris pH 7.5, 150 mM NaCl, 1 mM TCEP using a PD-10 desalting column and concentrated to ∼12 mg/ml. Crystals were grown in a 1:1 drop of protein to reservoir solution and equilibrated against 4.8 M ammonium acetate, 100 mM MES pH 5.5 at 4C. Crystals were cryoprotected by rapid transfer into reservoir solution supplemented with 25% glycerol before flash freezing in LN2. Diffraction data were collected on Advanced Light Source Beamline 5.0.2 and processed with XDS. The structure was determined by molecular replacement in Phaser using 6VO7 as a search model. The structure was refined using iterative rounds of refinement in REFMAC5 with manual inspection and model building in COOT. Ligand restraints for VVD-442 and the covalent bond with Cys242 were generated in JLigand. Waters were automatically added in COOT and REFMAC5 and manually inspected. Data collection and refinement statistics can be found in Supplementary Table 1.

### Intact protein mass spectrometry

To observe covalent ligand engagement of recombinant p110α, samples were analyzed following formic acid quenching on an Agilent LC1290 Infinity II instrument coupled to a 6545 QTOF liquid chromatography–mass spectrometer (Agilent Technologies). A sample volume of 10 μl, equivalent to approximately 1.5 pmol of p110α protein, was injected. The protein was desalted and separated on an AERIS 3.6-μM-wide-bore XB-C8 liquid chromatography column (50 × 2.1 mm^2^, Phenomenex) at 60 °C at a flow rate of 0.5 ml min^−1^. Liquid chromatography solvent A comprised 0.1% formic acid in 99.9% water, and solvent B was 0.1% formic acid in 99.9% acetonitrile. The column was equilibrated in 10% B for 30 s, followed by a 3.5 min gradient from 10 to 70% B to separate the analytes. This was followed by a 15 s gradient from 70 to 95% B, a 15 s gradient from 95 to 10% B, a 15 s gradient from 10 to 95% B and finally a 15 s gradient from 95 to 10% B to clean the column before re-equilibration. Mass spectra were acquired from 700 to 1,700 Da at a resolution of 25,000. A Dual Agilent Jet Stream Electrospray Ionization Source was used for ionization. The gas temperature was set to 325 °C, with a flow rate of 10 l min^−1^. The nebulizer was set to 45 pounds per square inch and sheath gas temperature and flow were set to 375 °C and 12 l min^−1^, respectively. One spectrum was acquired per second with a collision energy of 10 V. The capillary voltage was set to 5,000 V and the nozzle voltage to 2,000 V. The fragmenter, skimmer and octopole radio frequency (RF) peaks were set at 250, 65 and 750 V, respectively. The resulting data files were deconvoluted to protein masses using Agilent MassHunter BioConfirm Software, v.11.0. The biomolecule table containing protein mass and peak intensities was used to quantify the percentage of compound modification relative to the unmodified protein peak, by dividing modified protein intensity by the sum of the unmodified and modified protein intensities. *k*_obs_/[*I*] was calculated using assumptions for pseudo-first-order reaction kinetics (d(VVD-442)/d*t* = −*k* × (VVD-442), (VVD-442)*_t_* = (VVD-442)*_t_*_0_ × e^−*kt*^) and averaged for each inhibitor concentration; *k*_obs_ was determined by dividing these values by (*I*) for each concentration. *k*_obs_ values were plotted against inhibitor concentrations and the curve was fit with the equation *k*_obs_ =(k_inact_ *[VVD-442])/(K_i_ + [VVD-442])

### In vivo studies

#### Human cell-line derived xenograft (CDX) studies

For efficacy studies, following the acclimation period (3-7 days), immunodeficient mice, NSG (Jackson Laboratory) for A549 or NSG(Zhuhai BesTest Bio-Tech Co.,Ltd) for FaDu inoculation, and nu/nu(Jackson Laboratory) for 1975 and H2122 inoculation, were inoculated subcutaneously into the dorsal flank(A549, H1975, H2122, FaDu) or right shoulder(FaDu) with a cell suspension of tumor cells in 0.1 ml of PBS(FaDu, H2122, H1975) or a 1:1(v:v) mix of PBS and Geltrex(Cat #12760-021), delivering tumor cells (FaDu, H2122 and H1975 - 5 x 10^6^ cells/ mouse; A549 - 3X10^6^ cells/ mouse). Mice were randomized based on tumor volume and were enrolled into different groups. Tumor volumes were measured twice per week after randomization in two dimensions using a caliper.All treatment were administered at the indicated dose level by oral gavage. FaDu CDX TGI was conducted at Crown Biosciences.

For PK/PD/TE studies, following the acclimation period(3-7 days), immunodeficient NSG mice(Jackson Laboratory) were inoculated subcutaneously into the dorsal flank with a cell suspension of FaDu cells in 0.1 mL, 2 x 10^6^ FaDu cells/ mouse.

#### Patient derived xenograft (PDX studies)

PDX studies were conducted at Crown Biosciences(BR10564, GA0006, LU0876, CR2528, CR1554, CR0010 and LU5229) or Champions Oncology(CTG-3196 and CTG-3192). Fresh tumor tissues from mice bearing established primary human cancer PDX model was harvested and cut into small pieces (approximately 2-3 mm in diameter). PDX tumor fragment, harvested from donor mice, was inoculated subcutaneously at the upper right dorsal flank(Crown Biosciences) or left dorsal flank(Champions) into immunocompromised mice for tumor development. Mice were randomized based on tumor volume (100-200 mm3). After randomization, tumor bearing mice were allocated into indicated treatment groups, with 3 mice per group.

#### KPAR(^G12C^)

KRAS^G12C^ transplantation experiments were carried out in 8-10 week C57BL/6J mice. For subcutaneous experiments 150,000 cells were resuspended in PBS and mixed 1:1 with Geltrex LDEV-Free Reduced Growth Factor matrix (Thermo Scientific) and injected subcutaneously in one flank. For orthotopic lung tumors 150,000 KPAR^G12C^ cells were resuspended in 100 μl PBS and injected in the tail vein. Tumor volume was measured by micro-CT scan as previously described (DOI: 10.1038/s41596-022-00769-5)(*31*).

## References

1. S. R. Punekar, V. Velcheti, B. G. Neel, K. K. Wong, The current state of the art and future trends in RAS-targeted cancer therapies. Nat Rev Clin Oncol 19, 637–655 (2022).

2. P. Rodriguez-Viciana et al., Phosphatidylinositol-3-OH kinase as a direct target of Ras. Nature 370, 527–532 (1994).

3. P. H. Warne, P. R. Viciana, J. Downward, Direct interaction of Ras and the amino-terminal region of Raf-1 in vitro. Nature 364, 352–355 (1993).

4. J. A. Engelman et al., Effective use of PI3K and MEK inhibitors to treat mutant Kras G12D and PIK3CA H1047R murine lung cancers. Nat Med 14, 1351–1356 (2008).

5. M. Zhang, H. Jang, R. Nussinov, The structural basis for Ras activation of PI3Kα lipid kinase. Physical Chemistry Chemical Physics 21, 12021–12028 (2019).

6. S. Gupta et al., Binding of Ras to Phosphoinositide 3-Kinase p110α Is Required for Ras-Driven Tumorigenesis in Mice. Cell 129, 957–968 (2007).

7. E. Castellano et al., Requirement for Interaction of PI3-Kinase p110α with RAS in Lung Tumor Maintenance. Cancer Cell 24, 617–630 (2013).

8. S. E. Nunnery, I. A. Mayer, Management of toxicity to isoform α-specific PI3K inhibitors. Ann Oncol 30, x21–x26 (2019).

9. B. Vanhaesebroeck, M. W. D. Perry, J. R. Brown, F. André, K. Okkenhaug, PI3K inhibitors are finally coming of age. Nat Rev Drug Discov 20, 741–769 (2021).

10. F. André et al., Alpelisib for PIK3CA-Mutated, Hormone Receptor-Positive Advanced Breast Cancer. N Engl J Med 380, 1929–1940 (2019).

11. K. A. Baltgalvis et al., Chemoproteomic discovery of a covalent allosteric inhibitor of WRN helicase. Nature 629, 435–442 (2024).

12. M. E. Pacold et al., Crystal structure and functional analysis of Ras binding to its effector phosphoinositide 3-kinase gamma. Cell 103, 931–943 (2000).

13. R. Cooley et al., Development of a cell-free split-luciferase biochemical assay as a tool for screening for inhibitors of challenging protein-protein interaction targets. Wellcome Open Res 5, 20 (2020).

14. N. G. Martinez et al., Biophysical and Structural Characterization of Novel RAS-Binding Domains (RBDs) of PI3Kα and PI3Kγ. J Mol Biol 433, 166838 (2021).

15. M. Molina-Arcas et al., Development of combination therapies to maximize the impact of KRAS-G12C inhibitors in lung cancer. Sci Transl Med 11, (2019).

16. M. M. Murillo et al., Disruption of the Interaction of RAS with PI 3-Kinase Induces Regression of EGFR-Mutant-Driven Lung Cancer. Cell Rep 25, 3545–3553.e3542 (2018).

17. S. R. Rangan, A new human cell line (FaDu) from a hypopharyngeal carcinoma. Cancer 29, 117–121 (1972).

18. D. Goren et al., Targeting of stealth liposomes to erbB-2 (Her/2) receptor: in vitro and in vivo studies. Br J Cancer 74, 1749–1756 (1996).

19. J. Jiang et al., Translational and Therapeutic Evaluation of RAS-GTP Inhibition by RMC-6236 in RAS-Driven Cancers. Cancer Discov 14, 994–1017 (2024).

20. P. A. Jänne et al., Selumetinib Plus Docetaxel Compared With Docetaxel Alone and Progression-Free Survival in Patients With KRAS-Mutant Advanced Non-Small Cell Lung Cancer: The SELECT-1 Randomized Clinical Trial. Jama 317, 1844–1853 (2017).

21. A. J. de Langen et al., Sotorasib versus docetaxel for previously treated non-small-cell lung cancer with KRAS(G12C) mutation: a randomised, open-label, phase 3 trial. Lancet 401, 733–746 (2023).

22. M. M. Awad et al., Acquired Resistance to KRAS(G12C) Inhibition in Cancer. N Engl J Med 384, 2382–2393 (2021).

23. J. Canon et al., The clinical KRAS(G12C) inhibitor AMG 510 drives anti-tumour immunity. Nature 575, 217–223 (2019).

24. M. B. Ryan et al., KRAS(G12C)-independent feedback activation of wild-type RAS constrains KRAS(G12C) inhibitor efficacy. Cell Rep 39, 110993 (2022).

25. J. Boumelha et al., An Immunogenic Model of KRAS-Mutant Lung Cancer Enables Evaluation of Targeted Therapy and Immunotherapy Combinations. Cancer Res 82, 3435–3448 (2022).

26. S. Suire et al., Gbetagammas and the Ras binding domain of p110gamma are both important regulators of PI(3)Kgamma signalling in neutrophils. Nat Cell Biol 8, 1303–1309 (2006).

27. P. Rodriguez-Viciana, P. H. Warne, B. Vanhaesebroeck, M. D. Waterfield, J. Downward, Activation of phosphoinositide 3-kinase by interaction with Ras and by point mutation. Embo j 15, 2442–2451 (1996).

28. G. Q. Gong et al., A small-molecule PI3Kα activator for cardioprotection and neuroregeneration. Nature 618, 159–168 (2023).

29. D. Prakoso et al., Gene therapy targeting cardiac phosphoinositide 3-kinase (p110α) attenuates cardiac remodeling in type 2 diabetes. Am J Physiol Heart Circ Physiol 318, H840–h852 (2020).

30. J. Boumela et al., An Immunogenic Model of KRAS-Mutant Lung Cancer Enables Evaluation of Targeted Therapy and Immunotherapy Combinations. Cancer Research 82, 3435–48 (2022).

31. M. Zaw Thin et al., Micro-CT acquisition and image processing to track and characterize pulmonary nodules in mice. Nature Protocols 18, 990–1015 (2023).

32. M. Kavanagh et al. Selective inhibitors of JAK1 targeting an isoform-restricted allosteric cysteine. Nat Chem Biol. 18(12):1388–1398(2022).

